# Terminal maturation of human reticulocytes to red blood cells by extensive remodelling and progressive liquid ordering of membrane lipids

**DOI:** 10.1101/2023.06.02.543386

**Authors:** Giampaolo Minetti, Lars Kaestner, Harald Köfeler, Cesare Perotti, Isabel Dorn

## Abstract

In the age of “omics”, lipidomics of erythropoiesis is still missing. How reticulocytes mature in the circulation into functional erythrocytes is also for the most part unknown, beyond the lipidomics level. We have characterized here the lipid content of two subpopulations of peripheral reticulocytes of different maturity and three of erythrocytes of different age. Reticulocytes undergo profound changes in membrane lipid composition as they mature in the vasculature. Sphingomyelin and cholesterol increase, whereas phosphatidylcholine and phosphatidylserine decrease, relative to total lipids, from young reticulocytes to mature erythrocytes, suggesting that the area of the membrane in liquid-ordered state increases. The relative amounts of the more than 70 phospholipid subclasses evaluated here also change in the process. As peripheral reticulocytes and erythrocytes are unable of de-novo phospholipid synthesis, such remodeling likely requires selective removal of phospholipids from the membrane or their exchange with plasma or both. It has to be investigated whether this process might involve lipid transfer proteins that, when defective, such as in neuroacantocytosis syndromes, result in altered erythrocyte morphology. These findings not only shed light on fundamental aspects of red blood cell physiology and erythropoiesis but also raise intriguing questions surrounding protein-lipid interactions, membrane architecture, and lipid trafficking mechanisms.

## INTRODUCTION

The process of terminal differentiation from reticulocytes to mature red blood cells (RBCs) in mammals has long been a subject of intense scrutiny since the inception of RBC research^1^. Recent years have seen a resurgence of interest in this area^2–7^, particularly in light of the potential for culturing human RBCs *in vitro* for transfusion purposes^8–13^. This interest has been further fuelled by ‘omics’ approaches, which aim to reassess the properties of these cells in both health and disease^14–21^. While considerable knowledge has been amassed regarding the early phases of reticulocyte maturation within the bone marrow of adult mammals following enucleation of orthochromatic erythroblasts, the terminal development of circulating reticulocytes into mature RBCs remains elusive. Two primary challenges have impeded progress in this field: firstly, the scarcity of synchronized reticulocytes at a specific stage of differentiation. Studies have predominantly focused on “shift” reticulocytes^22^, obtained from humans with natural or induced reticulocytosis, or on “stress” reticulocytes^23^, after boosting reticulocyte counts in animals up to 90% of total red cells by phlebotomy or pharmacological treatment^24,25^. While synchronized to some extent, these cells may not accurately represent normal circulating reticulocytes^26,27^. Gradient centrifugation methods for reticulocyte enrichment, though yielding more “normal” reticulocytes, offer limited purification, thereby hindering precise biochemical characterization^28^. Additionally, cord blood provides a source of reticulocytes, albeit distinct from their adult counterparts^29,30^. Finally, non-nucleated cells obtained with variable efficiencies by culturing erythroid precursors *in vitro* are immature reticulocytes, probably akin to bone marrow reticulocytes at a very early stage of maturation. As a result, the issue as to whether peripheral circulating reticulocytes could mature into “normal” RBCs *in vitro* is controversial^31^. Secondly, characterization of terminal erythroid differentiation has been more recently conducted primarily through proteomics and transcriptomics approaches^2,32–37^, with limited attention to lipids as essential membrane constituents^34^. This occurred despite the fact that characterization of RBC lipids predated that of proteins, leading to seminal discoveries on membrane architecture in eukaryotic cells, such as the asymmetric transverse localization phospholipids in the two leaflets of the bilayer^38–41^. To address these challenges, we isolated subpopulations of circulating reticulocytes at various maturation stages *in vivo*, along with subpopulations of RBCs of different age from normal human donors and characterized their lipid content. Furthermore, we conducted a semi-quantitative analysis of protein VPS13A in all the cell subpopulations under study.

## RESULTS

### Cell size and shape

We have isolated, starting from a single sample of blood, two populations of CD71^+^ reticulocytes, RY and RM, representative of ≈0.1-0.2% younger and ≈0.4-0.5% more mature circulating reticulocytes. Differences in the maturation stage were assessed by analysing the relative differences in CD71 expression: the levels of CD71 in RY were 20 to 30 times the levels in RM (**Figure 1A**). Furthermore, the two populations of reticulocytes were free of contaminating mature RBCs (**Figure 1B**). Additionally, from the reticulocyte-depleted population of RBCs we have separated subpopulations of young (EY), middle-age (EM) and old (EO) erythrocytes, amounting, respectively, to the following percentages with respect to total cells subjected to density gradient separation: 5.2 ± 2.6, 93.3 ± 2.4 and 1.6 ± 0.3 (mean ± S.D.; n=4; **Figure 1C**). Analysis of the protein 4.1a/4.1b ratio confirmed that the subpopulations of RBCs of different density also differed by cell age (**Figure 1D**). EY may be considered as RBCs in their first week of life, during which they have matured from reticulocytes, whereas EO in their last few days of life, after 17 weeks in the circulation (see also Supplementary results and **Figure S1**). When examined by scanning electron microscopy, almost all reticulocytes displayed the morphology of a biconcave discocyte, with a few misshaped cells, mostly present in the RY samples (**Figure 2A-C and Figure S2**), indicating that reticulocytes are almost indistinguishable from mature RBCs from the morphological point of view at stasis. A progressive decrease in cell size from RY to RM and from RM to EM could be evaluated by measuring cell diameters, by bright field microscopy, in a sample of fixed cells. The projected area diameter of RY decreases by approximately ≈7.7% as the cells mature to EM (**Figure 2D**).

**Figure 1.**
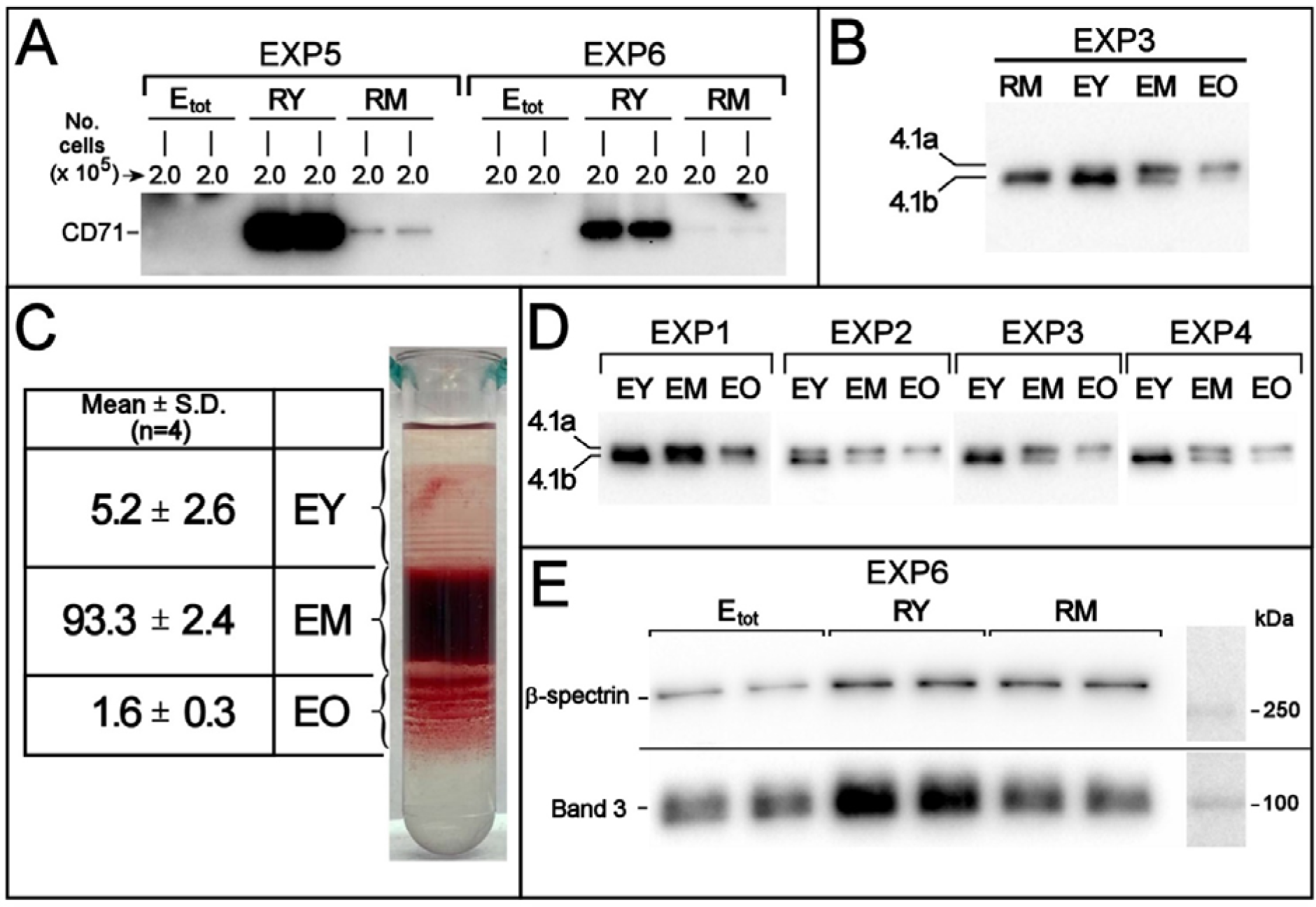
**(A)** Western blotting with anti CD71 of RY and RM reticulocytes and of total RBCs (E_tot_) from the corresponding donor in two representative experiments. For each sample, 2.0 x 10^5^ cells were loaded. The large differences in signal intensity between samples containing identical numbers of cells prevented quantification by densitometry of the bands. Instead, the relative differences in CD71 expression were evaluated by the procedure described in detail in the Supplementary material and **Figure S1A-C**. **(B)** Protein 4.1R Western blotting in reticulocytes and RBCs of different age. The blotting is representative of four independent experiments with identical results. RM, EY, EM and EO were from the same donor. In each lane, 2.0×10^5^ cells were loaded. See Supplementary material and **Figure S1D**. **(C)** Separation of RBCs according to density in self-forming Percoll^®^ gradients. One sample is shown as representative of four independent experiments. The table gives the average cell recovery in the three RBC subpopulations, evaluated as % Hb (or number of cells) recovered in each fraction with respect to total Hb (or number of cells) loaded in the gradient. **(D)** Western blotting of protein 4.1 in density-separated RBC subpopulations. The relative increase in protein 4.1a/4.1b ratio from young to old RBCs confirms that density separation resulted in a separation of RBCs of different age. Each sample contained 2.0 x 10^5^ cells. **(E)** Band 3 and β-spectrin levels in RY, RM and E_tot_ evaluated by Western blotting. The same number of cells (2×10^5^) was loaded for each sample, in duplicate. The blot shown is representative of four independent experiments with similar results. See also Supplementary material and **Figure S1G**.

**Figure 2.**
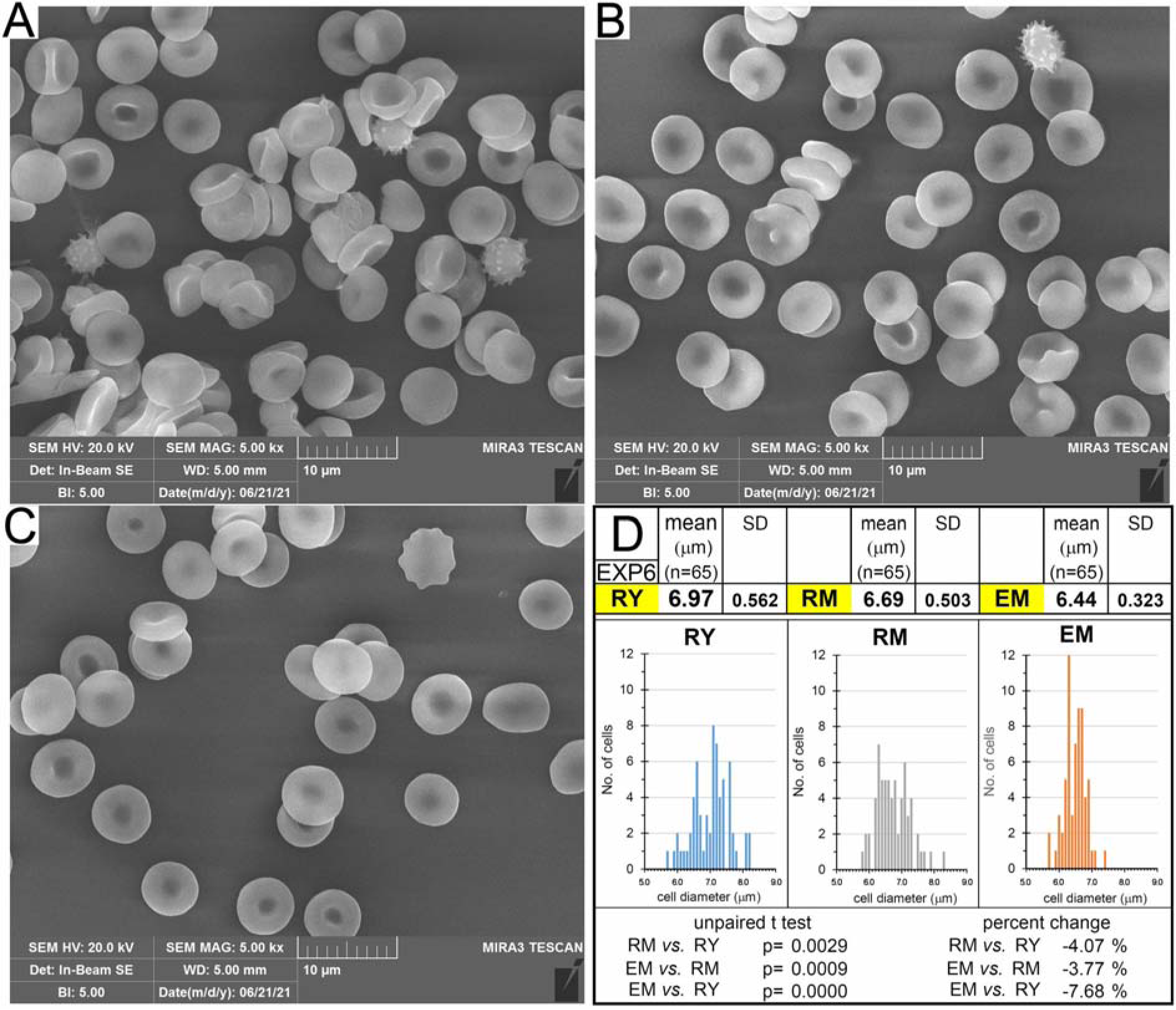
Scanning electron microscopy of **(A)** young circulating reticulocytes, RY; **(B)** mature reticulocytes, RM; **(C)** RBCs from the total population of reticulocyte-depleted cell sample of the same donor, E_tot_. Images are representative of four independent experiments with similar results. **(D)** Diameter of RY, RM and EM. Unstained cells were fixed with glutaraldehyde^57^, visualized in bright-field microscopy at 40X magnification and photographed with a digital camera. Thanks to a superimposed ocular micrometer, cell diameter were manually measured in the digital image. The histograms show the distribution frequencies of cell diameters in the three cell samples. The decreases in cell diameter were statistically significant to the unpaired Student’s t test. Data are representative of one of four experiments with similar results.

As RBCs age in circulation, they undergo coordinated membrane area and volume reduction to maintain a constant surface-to-volume ratio and prevent premature clearance due to spherization with consequent loss of deformability^42–46^. It is thought that the cell-age related membrane loss occurs through the spontaneous release of vesicles akin to those that can be obtained *in vitro* by stimulating RBCs with various manoeuvres, and that are devoid of membrane skeleton^47–49^. On the other hand, loss of membrane by early reticulocytes maturing in the bone marrow was described to occur via the release of exosomes^50^ enriched in transferrin receptor (TfR, CD71)^51,52^, and free of Band 3, glycophorin A and spectrin^53,54^. However, our research has demonstrated that human RBCs undergo a continuous loss of Band 3 and spectrin throughout their 120-day circulatory lifespan^55,56^. This led us to question whether the reduction in membrane area in circulating reticulocytes is accompanied by a loss of Band 3 and spectrin.

We observed a decrease in the abundance of these two proteins during the maturation of circulating reticulocytes (**Figure 1E**). We then posed a similar question regarding the lipid component. Membrane rafts, enriched in specific raft proteins and characterized by a cholesterol/sphingomyelin-rich liquid-ordered state^58^, have been described to be selectively lost with the exosomes released by reticulocytes in rats^59^. Consequently, one would expect the membrane of reticulocytes to become increasingly depleted in membrane rafts if the mechanisms that determine membrane loss were the same for bone marrow and circulating reticulocytes. As shown below, however, sphingolipids and cholesterol appear to be selectively retained in the membrane of reticulocytes maturing in the circulation, at the same time as their surface area decreases, suggesting a significant difference between the bone marrow and circulatory phases of reticulocyte maturation^60^.

### Lipids

The availability of subpopulations of reticulocytes and RBCs of defined cell age allowed a regression analysis of the changes in lipid content versus time over the entire life span of the human RBC. **Figure 3** shows the content of seven membrane lipids, as mol % of each species over total sample lipids, in the five cell populations under study. A relative increase in Chol and SM and a decrease in PC and PS from reticulocytes to RBCs is observed, confirming what previously reported by us when comparing a single population of reticulocytes with the total population of RBCs^61^. Starting from young reticulocytes (RY) the statistically significant relative increase in SM appears to plateau at the mature RBC stage (EM) (**Figure 3A**). The opposite behaviour is displayed by PC, with a significant decrease from RY to EM. PC also displays an increase phase in the second half of RBC circulatory life, i.e. from EM to EO (**Figure 3B**).

**Figure 3.**
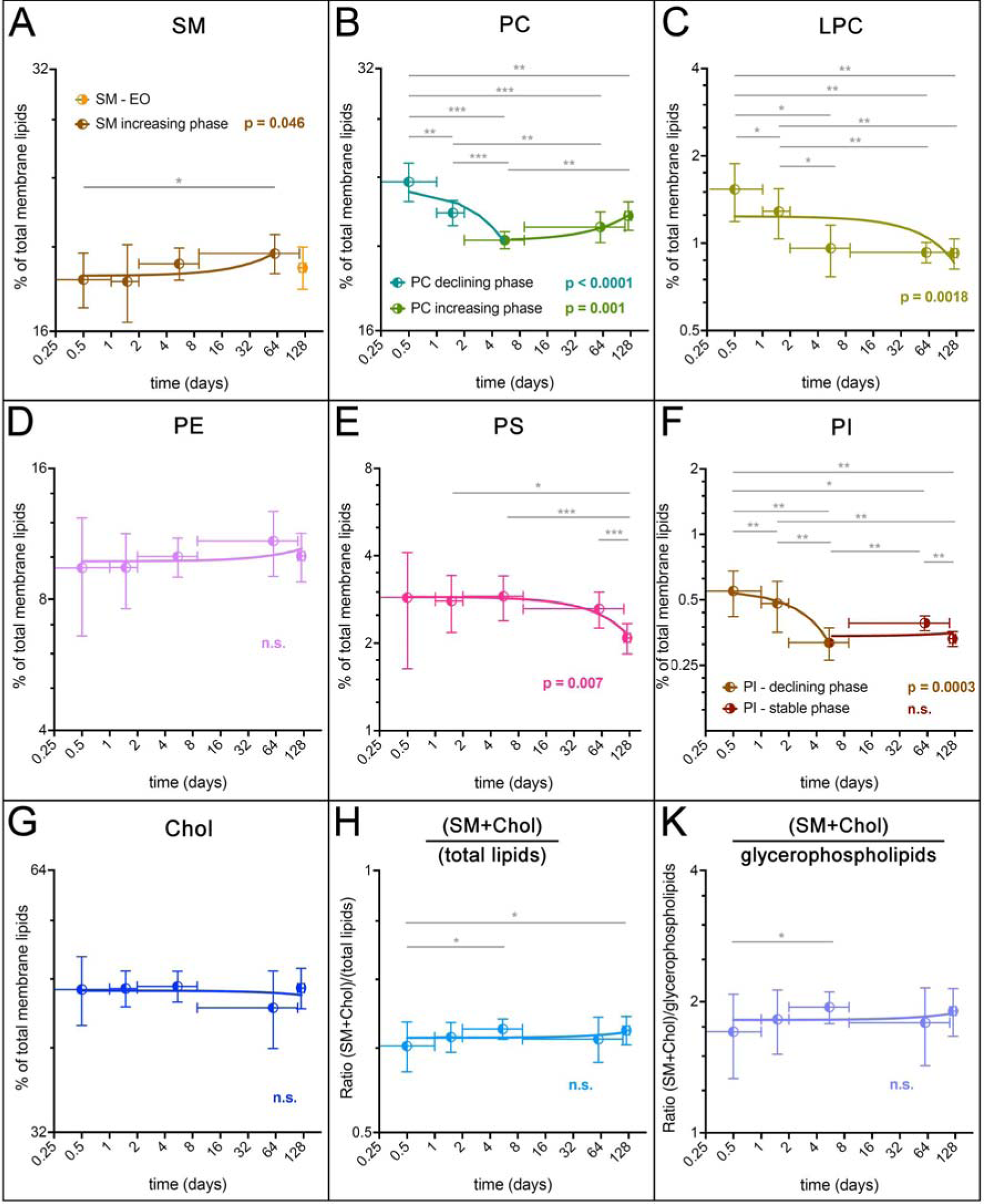
Regression analysis of the changes in membrane lipid content during maturation of circulating reticulocytes and aging of RBCs. The original values, expressed as percent of a given lipid with respect to total lipids from the same sample, have been logarithmically transformed and plotted against the time, in days, also on a logarithmic axis. Data were interpolated by linear regression and the statistical significance of the slope being different from zero, i.e. indicating a change, is shown in colour. Statistical analysis of pairs of samples is also shown, when significant, as horizontal lines connecting the two samples under test. N= 8. *: p < 0.05; ** p < 0.001; *** p < 0.0001. The tabulated original values and their direct graphical rendering are shown in **Figure S3** and **Figure S4**, respectively.

PS displays a continuous significant decrease all along maturation and ageing to the stage of old RBC (**Figure 3E**). LPC (**Figure 3C**) and PI (**Figure 3F**) are the other two phospholipids that, together with PS, display a significant decrease all along reticulocyte maturation and RBC aging. Concerning PE (**Figure 3D**) it displays a relative increase from the reticulocyte stage to EM, followed by a decrease in EO. However, these changes do not reach statistical significance (see also **Figure S4**). Concerning cholesterol, although its changes do not display statistical significance in any direction (contrary to the relative increase over total lipids documented in our previously published article^61^), when Chol and SM are together related to total lipids (**Figure 3H**) or to total glycerophospholipids (**Figure 3K**), a significant increase is again observed during maturation from reticulocyte to RBC (see also **Figure S4**).

### Phospholipid subclasses

When the different subclasses of a given phospholipid are analysed, they are seen to change, to variable extents, in relative abundance in the cell populations examined. Their changes should always be read in context with the underlying change in the relative amounts of that same phospholipid with respect to total lipids, be it a decrease or an increase (**Figure 3**). **Phosphatidylcholine. Figure 4** shows such changes for PC subclasses (for the corresponding tabulated values see **Figures S5** and **S6**). First, it is of importance to notice that the PC composition of RBCs taken from the literature^62^ (“E_tot_” in **Figure 4**) is practically superimposable to that of our sample of middle-age RBCs (EM). Our results show for the first time that analysis of phospholipid subclasses in the total population of RBCs is the average of values that differ significantly depending on the age of the cell. A continuous remodelling of phospholipid subclasses occurs all along the maturation from RY to RM, then EY and during the maturation and ageing of RBCs. In the case of PC, that shows an initial decrease relative to total lipids and then an increase from EM to EO (**Figure 3B**), individual PC subclasses change their relative amounts with a bimodal behaviour, all the way from RY to EO (**Figure 4**). One group of PC subclasses, containing saturated acyl chains, decrease relative to total PC [dipalmitoyl-PC (16:0/16:0), 1-palmitoyl,2-stearoyl-PC (16:0/18:0) and 1-palmitoyl,2-oleoyl-PC (16:0/18:1)], while and a second group, containing unsaturated acyl chains, displays a relative increase. In **Figure 4**, the composition in PC subclasses of plasma lipoproteins, (“PLASMA”) and total RBCs (E_tot_), taken from a literature dataset is also shown^62,63^. Interestingly, as RY mature to EM, the relative amounts of a given PC subclass (with a few exceptions) appear to approach the content of that particular subclass in plasma (see Discussion).

**Figure 4.**
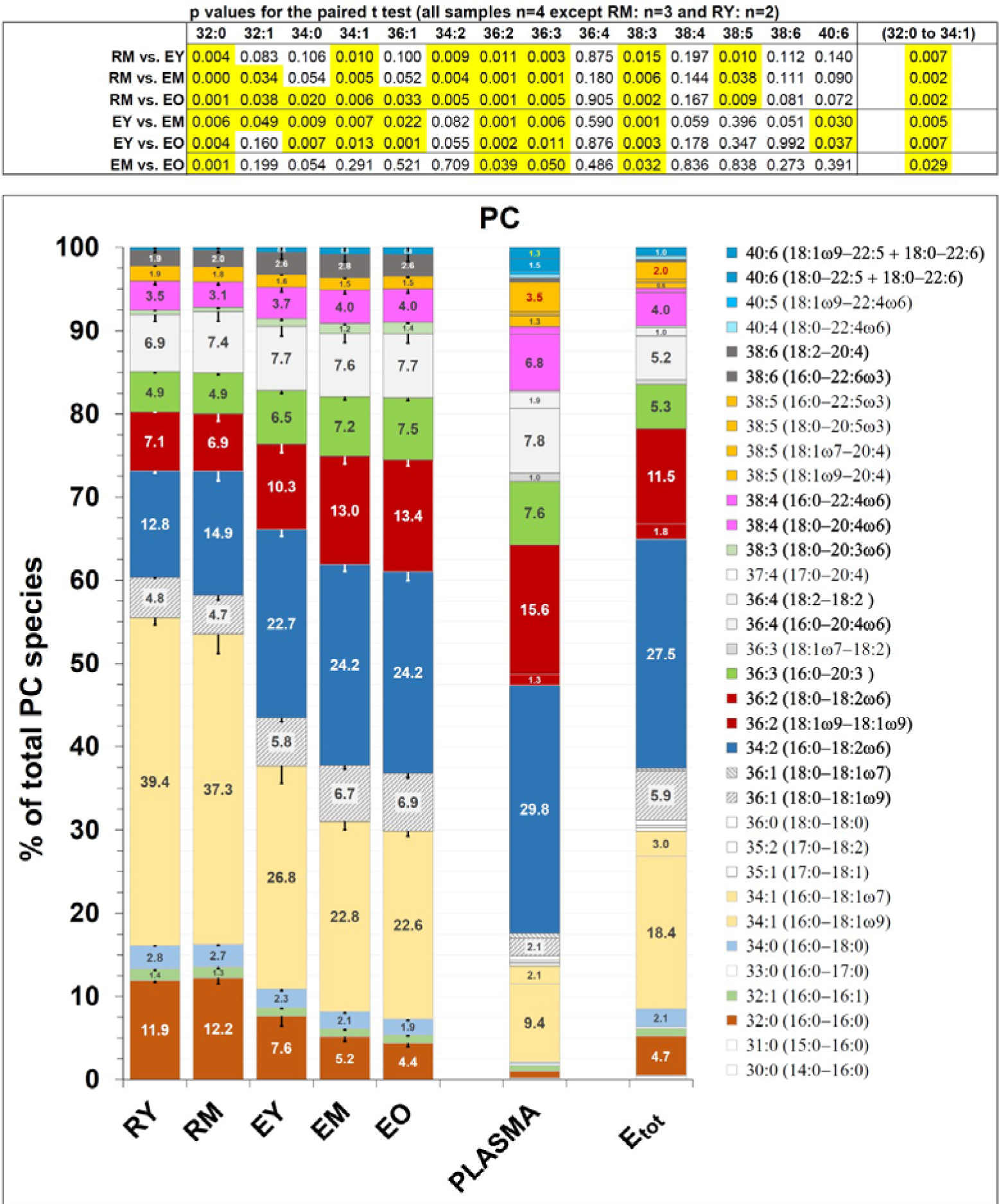
Diacyl PC subclasses in reticulocytes and RBCs. PC subclasses, as mol % of total PC in a given sample (as indicated by numbers inside the histogram bars) are presented with arbitrary colours that will be used consistently for the corresponding phospholipid subclass of diacyl PE, when present (Figure 5). The PC composition of samples “PLASMA” and “E_tot_” is taken from^62,63^, which was also used as a source for annotating the lipid subclasses obtained in our analysis. PC is ≈70% by weight of total lipids in plasma lipoproteins and ≈25 mole % of total phospholipids in RBCs^62,64^. The PC composition of EM is virtually superimposable to that of the literature sample Etot^63^. Moreover, the PC subclasses composition of plasma and RBCs was already described in earlier studies^65^, and it has been reported in a recent one^66^ with virtually identical results (see also **Figure S6**). The single saturated species (18:0/18:1) displaying a relative increase, instead of a decrease like all other saturated species (in bold in the legend to the right), is filled with dashed diagonal lines. Phospholipid subclasses were analysed in detail in a subset of 4 different donors. Error bars represent the standard deviation except for RY, where only two samples were available for analysis; here error bars represent the difference between the mean and the individual values. The table on top of the graph reports the results of the paired Student’s t test of comparisons among reticulocyte and RBC samples. Highlighted in yellow are the p values when ≤0.05. See also **Figures S5 and S6**.

#### Phosphatidylethanolamine

Diacyl PE subclasses also appear to change according to a pattern during the maturation of reticulocytes to RBCs. Again, this phospholipid’s subspecies can be divided in two groups. Those belonging to the first group display a relative increase with maturation, and contain relatively shorter and more saturated fatty acids (from 26:0 to 36:4). Those belonging to the second group contain longer and more unsaturated fatty acyl chains (from 38:4 to 40:7) (**Figure 5**, the corresponding tabulated values are shown in **Figure S7**). Changes appear specular to those undergone by PC subclasses but, unlike for PC, PE subclasses do not give the impression of a tendency to equilibrate with the corresponding plasma diacyl PE species, as changes go in the opposite direction (with a few exceptions) with respect to the PE plasma composition (see Discussion).

**Figure 5.**
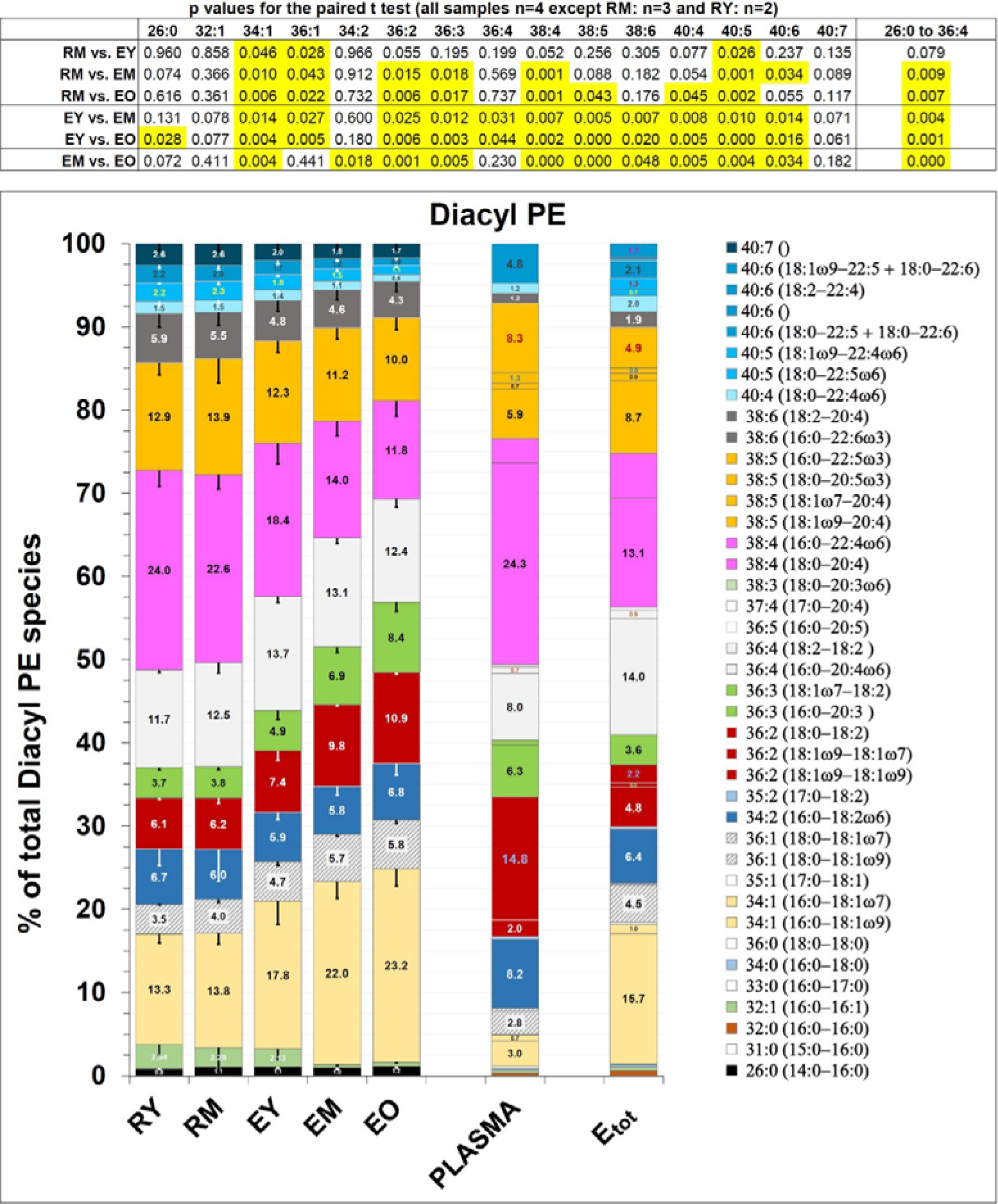
Subclasses of diacyl PE in reticulocytes and RBCs. Each subclass is given as mol % of total diacyl PE in a given sample. The diacyl PE composition of samples “PLASMA” and “E_tot_” is taken from^62,63^, which was also used as a source for annotating the diacyl PE lipid subclasses obtained in our lipidomics analysis (see Materials and Methods). Diacyl PE subclasses of corresponding fatty acid composition as PC subclasses shown in Figure 4 are shown in the same colour. Tabulated values can be found in **Figure S7**. For error bars and statistical analysis, see the legend to the previous Figure.

#### Sphingomyelin

Changes in the relative abundance of various SM subclasses occur with reticulocyte maturation, although to a lesser extent than for PC, and continue to the stage of EO (**Figure 6**) (tabulated values in **Figures S8-S10**). In **Figure 6**, the SM subspecies that are found in individual classes of plasma lipoproteins are also shown, taken from the literature^62,63^. Like for PC, most of the SM subclasses of the reticulocyte membrane give the impression of approaching the SM composition of plasma lipoproteins (except for the HDL family) as the reticulocyte matures. Furthermore, as with PC, a certain regularity of variation can be singled out. SM species carrying amidated fatty acids with less than 20 carbon atoms (from d32:1 to d36:2 included) display a relative increase in abundance, whereas those carrying a fatty acid with 20 or more carbon atoms decrease, pointing to a potential differential behaviour of inner *vs.* outer leaflet SM subclasses^67^.

**Figure 6.**
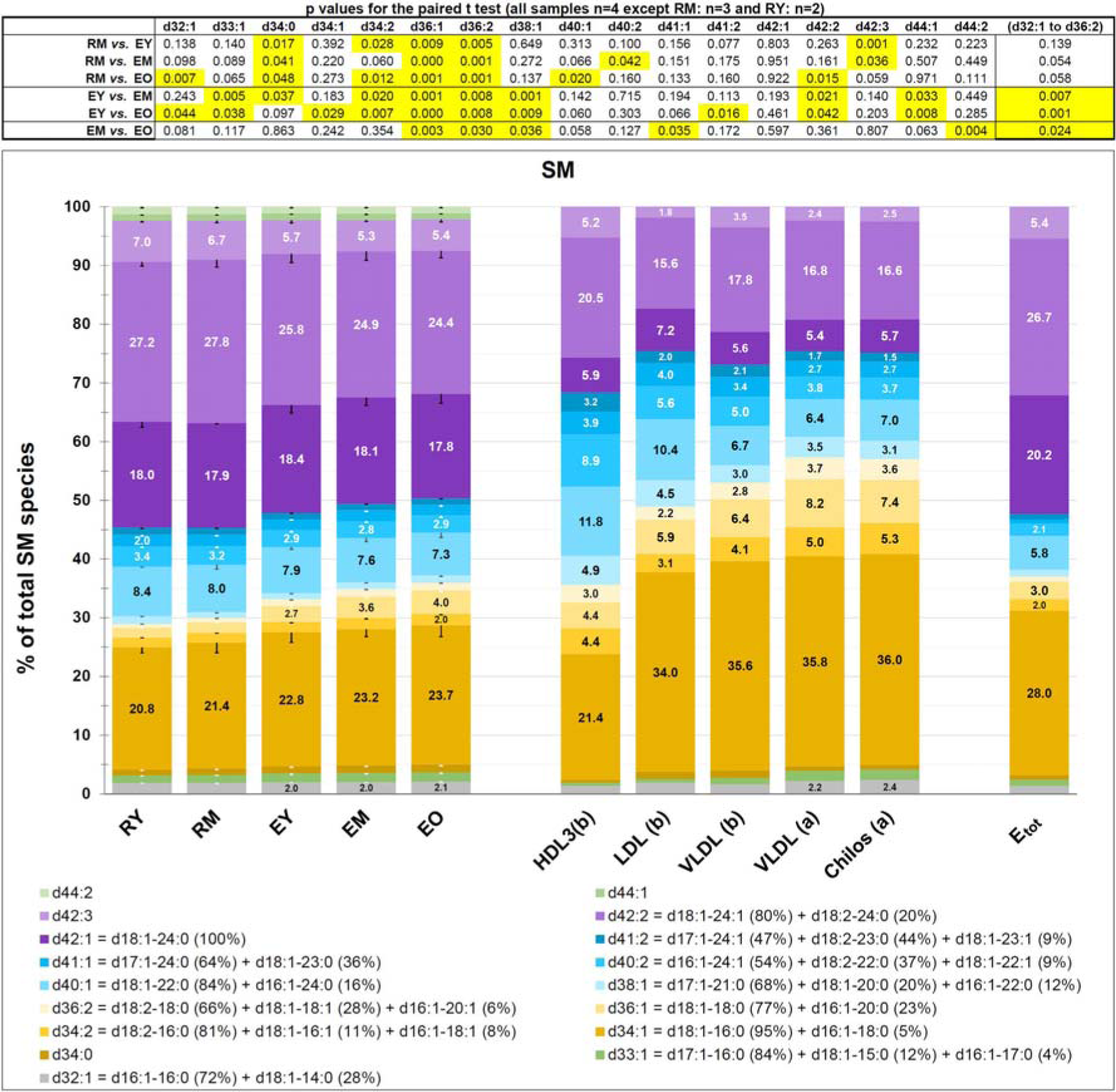
SM subclasses in reticulocytes and RBCs. Each subclass is given as mol % of total SM in a given sample, as indicated by numbers inside the histogram bars. The SM composition of “E_tot_” is taken from^62^, that of plasma lipoproteins from^63^. These two articles have been also used as a source for annotating the SM subclasses obtained in our lipidomics analysis. SM is ≈18% by weight of total lipids in plasma lipoproteins and ≈25% of total phospholipids in RBCs. Postprandial plasma of a normolipemic subject; (b) fasting plasma from three different normolipemic subjects^63^. For error bars and statistics, see previous Figure legend. See also **Figures S8-S10**.

#### Phosphatidylserine

PS subclasses with the longest and more unsaturated fatty acids decrease more, relative to total PS species, at the same time that total PS undergoes a decrease with respect to total membrane lipids, from RY to EO (**Figures 3E, S4 and S11**).

### Protein VPS13A in reticulocytes and RBCs

Chorein, now referred to as VPS13A, is a large protein found ubiquitously in normal tissues. Mutations in the *VPS13A* gene result in either the absence or significantly reduced expression of the protein in all tissues, including RBCs, in patients with Chorea acanthocytosis^68^. Further research revealed that VPS13A belongs to a large family of bridge-like lipid transfer proteins involved in non-vesicular lipid trafficking within eukaryotic cells. It facilitates lipid exchange at membrane contact sites between subcellular organelles and between organelles and the plasma membrane^69^. In Chorea acanthocytosis, loss of VPS13A function not only contributes to the neurological symptoms of the syndrome but also leads to abnormal RBC morphology. This abnormality may arise early in erythropoiesis due to impaired lipid trafficking in nucleated erythroid precursors and persist as a distinct feature of mature acanthocytes. Additionally, VPS13A might play a crucial role in lipid remodelling also in circulating reticulocytes and RBCs, where it is allegedly still present at high levels. However, because expression of VPS13A in mature RBCs was documented by immunoblotting of samples of whole blood^68,70^, concerns could be raised about possible artifactual results due to expression of the protein in white cells and/or platelets, rather than RBCs^71^. To address this issue, we conducted Western blotting analysis of VPS13A in all cell types examined in this study, ensuring the absence of contaminating white cells.

Results shown in **Figure 7A** confirm the presence of VPS13A in human reticulocytes and RBCs, and excludes a contribution from contaminating white cells or platelets to the observed result. Although VPS13 displays a progressive and continuous decrease from the reticulocytes to old RBCs (17^th^ week), it remains in the membrane at significant levels at least until the mature RBC stage, contrary to CD71 that is selectively and completely lost within the first week of circulatory life of the cell.

**Figure 7.**
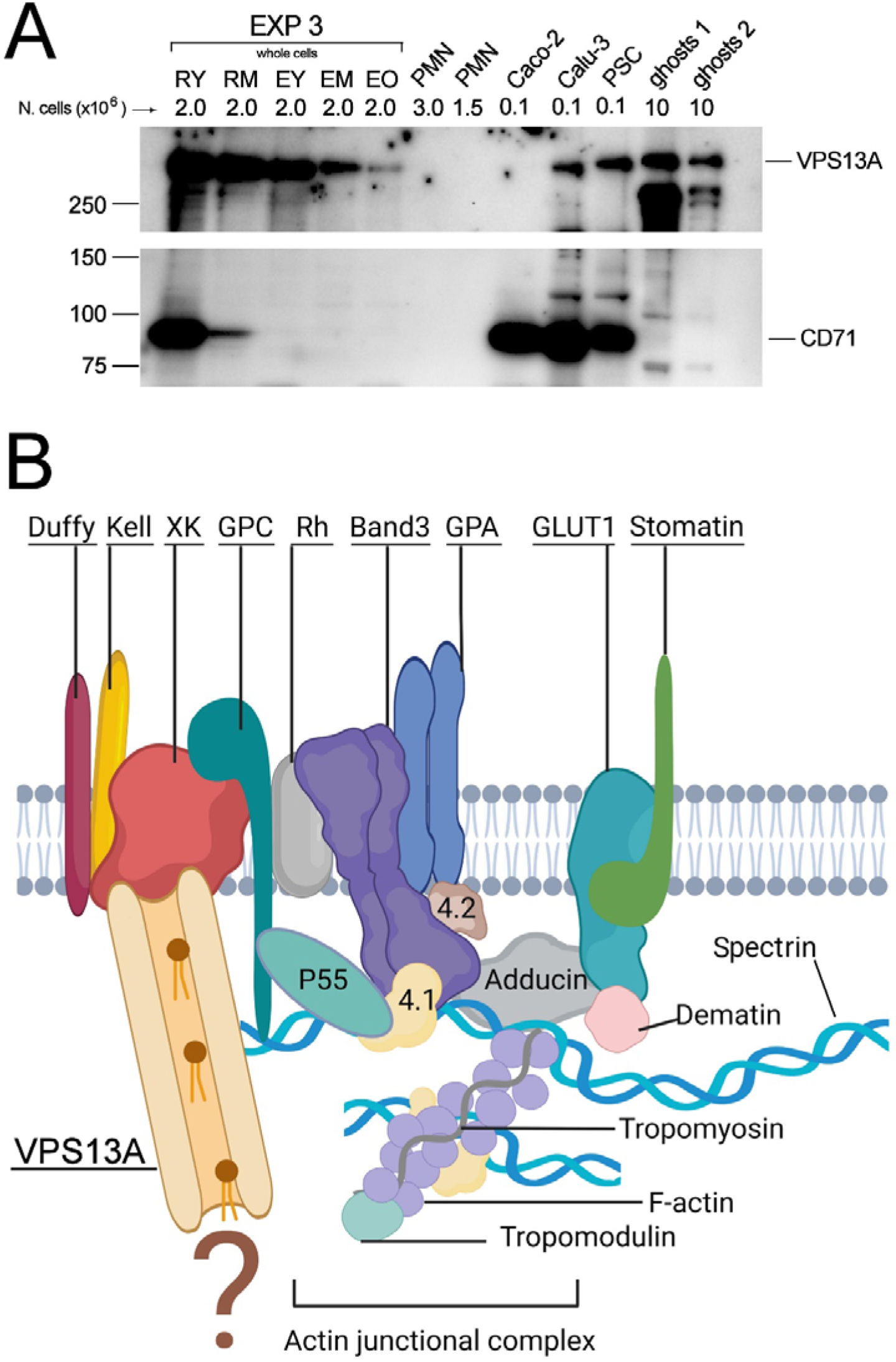
**A)** Western blotting of VPS13A in reticulocytes, RBCs and various other cell types. VPS13A was probed in reticulocytes and RBCs from one representative experiment, by loading equal amounts of whole cells dissolved in sample buffer, as indicated in the figure. Additionally, samples of polymorphonuclear leukocytes (PMN), Caco-2, Calu-3 and pancreatic stem cells (PSC), and purified ghost membranes from two different donors (ghosts 1 and ghosts 2), were loaded in the indicated amounts. The lower panel shows the same membrane re-probed for CD71. B) Schematic representation of the current model of the actin junctional complex in the RBC membrane^72^ and the possible disposition of VPS13A in the membrane as deduced from published data (see text for details).

## DISCUSSION

The heterogeneity of young red cells is a key aspect explored in this study. Unique to our research is the isolation of two distinct subpopulations of circulating reticulocytes: young RY and older RM, and three subpopulations of RBCs of different age: EY, EM, EO. Importantly, the isolated reticulocytes were “normal”, devoid of contamination by “stress” or “shift” reticulocytes^23^, as well as by mature RBCs, distinguishing them from previous studies. Based on arguments developed in detail in the Supplementary material, RY represent reticulocytes in their 12-24 hours post-egress from the bone marrow, while RM represent cells in the subsequent 24-48 hours of circulation. The subset of EY cells is indicative of RBCs in their first week of life. Conversely, EO cells correspond to RBCs in their 17^th^ week of life. The results of lipid analysis and morphological characterization allow reassessing some commonly accepted concepts about reticulocytes and their maturation. For instance, the timing for development of a mature RBC should be extended from the conventional 1-2 days in the circulation, to at least one week, as judged from lipid composition. Furthermore, our results support the frequently overlooked concept that circulating reticulocytes exhibit a biconcave discocyte morphology (**Figure 2**)^26,31,73^, contrary to the prevalent depiction of reticulocytes as roundish, irregularly shaped, multi-lobular cells^74^ (see also Supplementary material).

### Changes in lipid composition

Two main issues warrant discussion in light of the lipidomics data presented here. Firstly, understanding the mechanism(s) driving the selective remodelling of membrane lipids. Secondly, assessing the impact of these changes on RBC functionality. Considering the novelty of these findings in a relatively unexplored field, both issues will need additional research for full elucidation. Consequently, their discussion here may involve speculative considerations at times.

1. **Mechanisms responsible for membrane lipid remodelling**

We have previously observed, and confirm it here, that the lipid composition of mature RBCs displays a significantly higher ratio of (sphingolipids+Chol)/glycerophospholipids with respect to peripheral reticulocytes^61^. The observed changes can be most intuitively explained by the selective removal of certain lipids (PC, LPC, PI and PS), thereby leaving the remaining lipids either at constant levels (PE) or relatively enriched (SM and Chol) (**Figure 3**) in a membrane that undergoes continuous reduction in area throughout reticulocyte maturation and beyond^60^. In eukaryotic cells, phospholipids are distributed asymmetrically between the two leaflets of the plasma membrane. In human RBCs, the two abundant, choline-containing phospholipids, PC and SM, are both found prevalently (≈80%) in the extracellular leaflet. Among the aminophospholipids, virtually all PS and 80% of PE are located in the inner leaflet^39^. The membrane lipids undergo a remarkably intricate remodelling process, as demonstrated here, characterized by alterations in the relative proportions of no less than 70 distinct phospholipid subclasses throughout reticulocyte maturation. PC subspecies undergo selective substitution from the immature reticulocyte stage (RY) to young RBCs (EY), and from the latter to EM. To give only one example: the saturated species [dipalmitoyl-PC (16:0/16:0), 1-palmitoyl,2-stearoyl-PC (16:0/18:0) and 1-palmitoyl,2-oleoyl-PC (16:0/18:1)] decrease relative to total PC in a statistically significant manner from each of the cell subpopulations to the next more mature. In RY, these PC subclasses represent together ≈55% of total PC, whereas they are reduced to ≈30% in EM. The decrease in saturated PC species is mirrored by the relative increase of the species with longer and unsaturated acyl chains (**Figure 4**). Until now, mature RBCs have been defined based on the absence of any detectable remnants of TfR in the membrane and of RNA in the cytoplasm with respect to reticulocytes. A more accurate definition should now be based on the inversion of the ratio between the relative amounts of palmitoyl-oleoyl (16:0/18:1) and of palmitoyl-linoleoyl (16:0/18:2) species of PC, which changes from > 2 in reticulocytes to <1 in mature RBCs, making young RBCs more similar to reticulocytes than to mature RBCs (**Table S1**). Concerning PE (**Figure 5** and **S7**), the subspecies carrying more saturated acyl chains increase relative to total PE, whereas more unsaturated PE subclasses decrease, with the result that changes in PE fatty acid composition appear to coarsely mirror those of PC subspecies (**Figure 4**).

A mechanism such as the “pinching off” of membrane vesicles by macrophages, with recognition of membrane regions enriched in PC subclasses with more saturated acyl chains, which are those displaying a relative decrease during reticulocyte maturation (**Figure 4**), could explain the selective changes in subclass composition also for other phospholipids. Alongside these PC species, there must be equivalent amounts of an interdigitated inner-leaflet phospholipid, or a mixture of phospholipids, that are removed concurrently with PC. Among these species, PS would be present, as it decreases during reticulocyte maturation and RBC aging relative to total lipids (**Figure 3E**). If also some selected PE subclasses in the inner leaflet, enriched in unsaturated acyl chains, interdigitate with PC, the removal from the outside of the cell of more saturated PC species would result in their simultaneous extraction, thus accounting for the coarsely mirrored direction of changes in PC and PE subclasses (**Figures 4** and **5**). The fact that PE remains constant (although a statistically non-significant increase can be perceived) relative to total lipids during the transition from reticulocyte to mature RBC (**Figure 3D**) could be due to the predominant presence of PS compared to PE in the fraction of internal lipids co-extracted with PC.

According to the same mechanism, SM should be largely spared from the extraction from the membrane, resulting in the observed relative increase of SM over total phospholipids in mature RBCs with respect to reticulocytes (**Figure 3A**). However, some kind of selection may also affect SM, because SM species with less than 20 carbon atoms in their acyl chain display a relative increase over total SM from reticulocytes to RBCs and during RBC ageing (**Figures 6** and **S10**). Interestingly, ≈70% of these “shorter” SM species appear to be located in the inner leaflet, which contains ≈20% of total SM, while ≈70% of the longer species are in the outer leaflet, which contains the remaining ≈80% of SM^67^. It will require future investigation to understand the reason of this asymmetric remodelling of SM subspecies and while it continues as RBCs age in the circulation.

In addition to the mechanism outlined above, is important to acknowledge other processes that could contribute to the observed lipid remodelling, as previous literature has offered experimental evidence of their presence in mature RBCs. However, it should be noted that they have not been integrated into a comprehensive model associated with the maturation of reticulocytes. The lipid composition of the reticulocyte as it enters the circulation is the result of de novo lipid synthesis taking place during erythroid development and of possible lipid exchange with the extracellular milieu. Circulating reticulocytes have lost the internal membranes on which phospholipid metabolism takes place^60,61,75^. Indeed, none of the enzymes of the Kennedy’s pathway, responsible for the last steps in the biosynthesis of PC and PE in eukaryotic cells, including erythroid precursors^16^, have been detected by proteomic analysis of human peripheral reticulocytes^76^. However, reticulocytes can exchange of entire phospholipid molecules between the membrane and plasma^77–81^, the source of lipids being plasma lipoproteins. Moreover, some phospholipid classes can be translocated across the bilayer and remodelled in their acyl chains through the Lands cycle^82–84^. The exchange with plasma would involve primarily PC and SM, consistent with the phospholipid composition of plasma lipoproteins: ≈70% PC by weight, ≈20% SM and less than 5% for the other phospholipids^62,64^. Our data seem to indicate that an exchange of PC could be driven by the concentration gradient existing between cells and plasma for each of the PC subspecies. Thus, the relatively higher percent of polyunsaturated PC subspecies in plasma would drive an increase of these species in the reticulocyte, where they are at lower levels, to make them the most abundant species in the RBC, while the saturated species would move in the opposite direction (**Figure 4**). To account for changes in subclasses of phospholipids of the inner leaflet, which cannot directly exchange with plasma, alternative mechanisms need to be considered. For PE (**Figure 5**), the simplest possibility is that some selected PE subclasses are actually extracted from the outer leaflet in exchange with plasma PE, because, contrary to PS, which is entirely confined in the inner aspect of the bilayer, ≈20% of PE is located in the outer leaflet in mature human RBCs^39^. This is however unlikely, because of the extremely low levels of PE in plasma lipoproteins^63,64^ and the fact that changes in PE subclasses appear to go against a hypothetical equilibration with plasma (**Figure 5**). The exchange could be heterologous, between PE in the cell membrane and some other phospholipid from lipoproteins, or occur in a concerted manner with changes in PC, for instance through direct transfer of fatty acids from PC to PE^85^. Another potential scenario to consider is the net extraction from the membrane of specific individual phospholipid molecules without exchange with another phospholipid as it was described to occur in bovine RBCs^86^.

Amidst this uncertainty, what appears reasonably certain is that reticulocytes (and RBCs), lacking the capability for spontaneous de novo lipid biosynthesis and membrane trafficking^61^, must depend on external conditioning, not necessarily involving only splenic macrophages. In fact, other organs, such as the liver, have been shown to play an important role in processing RBCs for repairing and remodelling^87–90^. These agents and the mechanistic details by which they operate still have to be identified. With this, it will become clear if the active removal of areas of the membrane is sufficiently selective to produce the observed changes in phospholipid subspecies or exchange with plasma lipoproteins must also be invoked. Further investigations will also help understanding whether intracellular lipid transfer proteins such as VPS13A. This protein, formerly known as chorein, was found to be expressed in virtually all tissues examined, including RBCs^69,76^. We have confirmed its expression here and showed that although it declines significantly as reticulocytes and young erythrocytes mature, it is still present at significant levels in mature RBCs (**Figure 7A**). Figure 8B displays a hypothetical localization of VPS13A in the RBC membrane. Its interactions with membrane proteins have been described in the literature, in particular with XK^91,92^. Loss-of-function mutations in VPS13A have been associated with the acanthocyte phenotype of RBCs as observed in chorea-acanthocytosis^69^. It is intriguing that a protein facilitating lipid exchange at contact sites between membranes of intracellular organelles remains unaffected by the gradual breakdown of unnecessary components typical of reticulocyte maturation and it is found in significant quantities in mature RBCs, which lack internal membranes. One potential avenue for elucidating this phenomenon lies in exploring the interaction between VPS13A and the scramblase XK. Notably, XK is present in the mature RBC membrane and mutations in XK lead to the clinical manifestation of McLeod syndrome, characterized, like in chorea-acanthocytosis, by alterations in RBC morphology alongside neurological symptoms^93^. It is conceivable that VPS13A could supply the lipids necessary for XK-mediated scrambling between the two leaflets of the membrane, thereby facilitating coordinated changes in the composition of these leaflets. However, it remains a mystery how VPS13A could act according to this mechanism in a cell type that is lacking internal membranes from where to derive the required lipids (question mark in **Figure 7B**). It is therefore of utmost interest to investigate further on the possibility that VPS13A might play a role in RBC development^94^.

II) **Consequences of membrane lipid remodelling**

The functionality of protein-mediated membrane transport systems is known to be influenced by the lipid composition of the membrane, as indicated by previous studies^95–98^, and it is therefore anticipated that it will change along with reticulocyte maturation. For instance, the rate of Band 3-mediated anion exchange and of active K^+^ transport were found to be sensitive to the nature of the surrounding phospholipids and their fatty acid content^99,100^. Passive K^+^ transport, that was originally considered as the residual “leak” transport (although it was later attributed mostly to the K^+^(Na^+^)/H^+^ exchange^101^), was also found to correlate positively with the total content of arachidonic acid in the membrane phospholipids^101–103^. Fine-tuning of PIEZO1 function in RBCs depends on the fatty acid composition of membrane phospholipids^104^. Two important fatty acids, arachidonate (C20:4) and linoleate (C18:2) undergo significant changes during reticulocyte maturation and RBC ageing (Supplementary Material, **Figure S14**). Further studies will be required to shed light on this important issue. Due to the increased relative levels of Chol and SM in relation to total lipids, it is anticipated that the membrane of mature RBCs will exhibit a higher degree of liquid-ordered state compared to that of the reticulocyte^105^, potentially influencing membrane functionality. Membrane rafts, as regions of the plasma membrane in liquid ordered state, have been proposed to play important roles, from signal transduction, to cell-cell, cell-extracellular matrix and cytoskeleton-membrane interactions. We have shown for the first time that membrane rafts in RBCs are tightly associated with the spectrin skeleton through electrostatic interactions^105,106^ that do not involve the “canonical” Band 3-ankyrin and actin-protein 4.1-p55-GPC junctional complexes^72^. Therefore, we proposed that spectrin could interact directly with the polar head-groups of some subclasses of raft phospholipids. The literature previously described the affinity of spectrin for PS and PE, with subsequent research primarily emphasizing PS, while the significance of PE has been somewhat overlooked^107–111^. Yet, evidence showed that PE displays the highest affinity for spectrin^112,113^. It is remarkable that, in front of considerable variability, in RBCs from different mammals, in the relative amounts of other phospholipids, PE displays the lowest variability of all^95^ (**Table S2**). Taking a fresh perspective on the extensively studied topic of membrane skeleton biogenesis^114,115^, we suggest that the transformation of a relatively large reticulocyte (which turns out to be already a discocyte, as shown here) into a smaller biconcave discocyte is primarily driven by a controlled reduction in cell volume and the loss of specific bilayer regions, rather than solely by the assembly of the spectrin skeleton. The discoid shape does not inherently signify the presence of a fully assembled membrane skeleton^6,116,117^, as additional steps are required for its complete maturation from the already available full set of membrane skeletal components^118^. The relative instability of reticulocytes compared to mature RBCs^119–122^ can be attributed to the skeletal network still being somewhat “threadbare,” although it may already be “pinned” to the cytoplasmic leaflet by the interaction of spectrin with PE^123,124^. Early observations indicating that newly synthesized spectrin attaches to the plasma membrane through interactions distinct from spectrin-ankyrin and spectrin-protein 4.1, preceding its assembly into an extended meshwork^118,125–128^, align perfectly with this model. This unique positioning of spectrin results in a “reduction of dimensionality”^129^, thereby enhancing the rate of membrane skeleton assembly as the cell concurrently undergoes a reduction in volume and membrane area. The establishment of horizontal interactions and the anchoring to the bilayer via protein-protein vertical interactions could be considered complete only in mature RBCs^130^.

## CONCLUSIONS

The results obtained may have significant implications for several unresolved issues in RBC biology. The intricate process of lipid remodelling in reticulocytes preceding their transition into functional circulating RBCs warrants further exploration to uncover its underlying mechanisms. This inquiry holds broad significance, particularly in elucidating protein-lipid interactions pertinent to membrane transport systems and membrane skeleton biogenesis. Understanding the possible role of lipid transfer proteins in facilitating lipid exchange between reticulocytes/RBCs and their environment is crucial. Identifying the key regulators of reticulocyte maturation in the bloodstream holds promise for generating fully mature RBCs *in vitro* for transfusion purposes. While the findings presented here do not offer a precise sequence of events leading to the observed differences, they illuminate several potential avenues for future research in this domain.

## METHODS

A detailed version of Materials and Methods is available in the Supplementary material. Human blood samples were collected in lithium heparin as the anticoagulant at the local Transfusion center (Servizio di Immunoematologia e Medicina Trasfusionale of the IRCCS Policlinico San Matteo, Pavia, Italy) from regular donors after informed consent was obtained (protocol approved on 2017/04/10, by the local ethics committee: Comitato Etico Area Pavia, IRCCS Policlinico San Matteo, Pavia, Italy). Blood was leukodepleted and RBCs suspended at ≈20 % haematocrit (Ht) in PBSG (5 mM sodium phosphate pH 7.4, 154 mM NaCl, 4.5 mM KCl, 305-310 mosmol/kg H_2_O). Aliquots were used for cell count, Ht and Hb determination (Drabkin’s reagent), SDS-PAGE and Western blotting, fixation for SEM, and lipid extraction for lipidomics analysis. The RBC suspension was then used for the immunomagnetic isolation of CD71^+^ reticulocytes by using the “CD71 MicroBeads” system (Miltenyi Biotec, Germany). By modulating the flow rate of the RBC suspension passing through the magnetized separation columns, two different populations of reticulocytes were isolated in a number of independent experiments with blood from different donors, and named RY, for “young reticulocytes” and RM for more “mature reticulocytes”.

### Separation of RBCs into subpopulations of different age

After isolation of CD71^+^ reticulocytes, the reticulocyte-depleted RBC suspension from four different donors was subjected to centrifugation in self-forming Percoll^®^ Plus (GE Healthcare, Milan, Italy) gradients according to a previously published method^131^, with some modifications (see Supplementary material). Three subpopulations of cells of different density were isolated from the gradients: young erythrocytes (EY), erythrocytes of middle-age (EM) and old erythrocytes (EO).

### Preparation of cells for Scanning Electron Microscopy (SEM)

Before preparation for SEM, RBCs and reticulocytes were fixed with glutaraldehyde (GA) according to a two-step protocol^57^. Fifty μl of a suspension in water of GA-fixed cells were mixed with 50 μl of 2% osmium tetroxide in water (Cod. 75632-5ML, Merck Life Science S.r.l., Milan, Italy). After 30 min at room temperature, cells were washed with deionized water and were subjected to graded alcohol treatment, starting from 200 μl 70% ethanol, followed after 30 min by 80% ethanol and, after another 30 min, by absolute ethanol, in which cells could be stored at 4 °C. For SEM, a few μl of these suspensions were poured on the surface of a 5 mm x 7 mm silicon chip (Ted Pella Inc, Redding, CA, USA). After evaporation of the solvent, the chip was laid on one side of a bi-adhesive plastic disc coated with conductive material (Ted Pella Inc.) which was in turn attached to the surface of a standard pin stub mount for SEM. The surface of the samples was made electrically conductive by a coating of Pt using a Cressington 208HR Sputter Coater. SEM images were obtained through a FEG-SEM TESCAN Mira3 XMU, Variable Pressure Field Emission Scanning Electron Microscope (TESCAN, Brno, Czech Republic) located at the Arvedi Laboratory, CISRiC, Pavia, Italy. Observations were made, at different magnifications, in secondary electrons mode at 20 kV with an In-Beam SE detector at a working distance of 5 mm.

### Lipid analysis

Aliquots of packed reticulocytes or RBCs, containing 2×10^7^ cells, were extracted with 0.5 ml + 0.5 ml of ice-cold, HPLC-grade methanol and transferred to 10 ml Pyrex^®^ glass tubes with polyphenolic screw cap (1636/26MP, Italtrade S.r.l., Genoa, Italy). The quantitative lipid measurement was done by high-resolution mass spectrometry (LTQ-Orbitrap, Thermo Scientific)^132^. Lipids were extracted with an established MTBE (Methyl-tert-butylether) protocol^133^. The Orbitrap Velos Pro hybrid mass spectrometer was operated in Data Dependent Acquisition mode using a HESI II ion source. Full scan profile spectra from m/z 450 to 1050 for positive ion mode and from m/z 400 to 1000 for negative ion mode were acquired in the Orbitrap mass analyser at a resolution of 100k at m/z 400 and < 2 ppm mass accuracy. Samples were measured once in positive polarity and once in negative polarity. For MS/MS experiments, the 10 most abundant ions of the full scan spectrum were sequentially fragmented in the ion trap using He as collision gas (CID, Normalized Collision Energy: 50; Isolation Width: 1.5; Activation Q: 0.2; and Activation Time: 10). Centroided product spectra at a normal scan rate (33 kDa/s) were collected. The custom developed software tool Lipid Data Analyzer was used for data analysis^134,135^. Seven classes of lipids were quantified, together with their subclasses according to the length and number of double bonds of the acyl chains linked to the glycerol or sphingosine moiety: PC, lysophosphatidylcholine (LPC), SM, PE, PS, phosphatidylinositol (PI) and Chol. The relatively low amounts of reticulocytes that could be isolated in this study restricted the lipidomics characterization to direct MS analysis without intermediate chemical steps, providing only limited information on the chemical structure of each phospholipid species^136^. Yet, some of the missing information was recovered by annotating most possible phospholipid subclasses based on highly valuable, previously published characterizations of the human RBC lipidome^62,137^. Related Figures in the Supplementary material give additional details on the issue. Concerning PE, we have examined in detail the diacyl species, which amount to ≈49% of all PE, the rest being plasmalogens (alkylacyl ≈3%; alkenylacyl ≈48%) in human RBCs^62,63^. Although detailed characterization of PE plasmalogens in reticulocytes and RBCs is in progress, it can be for the moment assumed that diacyl and alkenylacyl PE species change in unison during reticulocyte maturation. All the more, since it is known that a correlation exists between the amounts of a given plasmenyl PE subspecies and the corresponding diacyl PE subspecies, indicating the common metabolic origin. Moreover major PE reorganization in the cell membrane due to exchange with plasma PE can be excluded because of the extremely low levels of PE in plasma lipoproteins^63,64^.

### SDS-PAGE and Western blotting

Standard protocols were adopted for SDS-PAGE in 10% isocratic or 5%-15% gradient polyacrylamide mini-gels (Bio-Rad Laboratories S.r.l., Segrate, Italy), and for Western blotting by electro-transferring proteins from gels to PVDF membranes (0.2 μm pores) using a Trans Blot Turbo system (Bio-Rad Laboratories S.r.l., Segrate, Italy) according to manufacturer’s instructions. After incubation with the primary antibody, and the appropriate secondary horseradish-peroxidase-(HRP)-conjugated antibody, membranes were developed with the chemiluminescence kit Prime Western Blotting Detection Reagent (GE Healthcare, Milan, Italy) and the signal was acquired with a Molecular Imager ChemiDoc XRS+ (Bio-Rad Laboratories S.r.l., Segrate, Italy). Densitometry of the bands was performed using the software Scion Image (Scion Corporation, USA). Antibodies usedwere: mouse monoclonal (BIII-136) anti human Band 3 (B9277, Merck Life Science S.r.l., Milan, Italy). Mouse monoclonal (H68.4) anti human CD71, sc-51829; mouse monoclonal anti human β-spectrin (VD4), sc-53901 (Santa Cruz Biotechnology, Dallas, Texas, USA); rabbit polyclonal anti human VPS13A (28618-1-AP, Proteintech Germany GmbH, Planegg-Martinsried, Germany). HRP-conjugated goat anti-mouse IgGs (170-6516) and HRP-conjugated goat anti-rabbit IgGs (170-6515) (Bio-Rad Laboratories S.r.l., Segrate, Italy).

## LIST OF ABBREVIATIONS

CD71, transferrin receptor; Chol, cholesterol; GPA, glycophorin A; LPC, lysophosphatidylcholine; PC, phosphatidylcholine; PE, phosphatidylethanolamine; PI, phosphatidylinositol; PS, phosphatidylserine; RBC, red blood cell; SM, sphingomyelin; Tfr, transferrin receptor; VPS13A, vacuolar protein sorting 13 homolog A.

## ACKNOWLEDGMENTS

Work supported by the EU Commission Horizon 2020 Marie Skłodowska-Curie Actions Innovative Training Networks project RELEVANCE Grant Agreement N. 675115 (to GM) and INNOVATION Grant Agreement N. 101120168 (to LK). Research conducted within the “Dipartimenti di Eccellenza” program (2018–2022) of the Italian Ministry of Education, University and Research (MIUR) - Dept. of Biology and Biotechnology “L. Spallanzani”, University of Pavia (GM).

## AUTHOR CONTRIBUTIONS

G.M. designed and conducted the experiments and wrote the paper. I.D. and H.K. performed analyses and wrote the paper. All Authors elaborated data and wrote the paper.

## SUPPLEMENTAL INFORMATION TITLES AND LEGENDS

Supplementary material: PDF file containing detailed methods, supplemental results in the form of figures and tables, and relevant references.

