## Supplementary material for "Terminal maturation of human reticulocytes to red blood cells by extensive remodelling and progressive liquid ordering of membrane lipids"

#### SUPPLEMENTARY MATERIALS AND METHODS

**Blood samples.** Human blood was collected in standard 6 ml vials containing lithium heparin as the anticoagulant by the local Transfusion center (Servizio di Immunoematologia e Trasfusione" of the IRCCS Policlinico San Matteo, Pavia, Italy) from regular donors after informed consent was obtained, according to the protocol approved by the local ethics committee (Comitato Etico Area Pavia, IRCCS Policlinico San Matteo, Pavia, Italy).

The blood was filtered to separate RBCs from white blood cells and platelets as previously described in detail[1]. The resulting purified RBCs were suspended at ≈20 hematocrit (Ht) (the actual value was measured using a micro-hematocrit centrifuge), in PBSG (5 mM sodium phosphate pH 7.4, 154 mM NaCl, 4.5 mM KCl, 305-310 mosmol/kg $H_2O$). Aliquots were used for cell count, Ht and Hb determination (Drabkin's reagent), SDS-PAGE and Western blotting, fixation for SEM and lipid extraction for lipidomics analysis.

**Immuno-magnetic separation of Retics**. All the procedures involving handling of reticulocytes were performed using buffers and materials at 4 °C. The purified RBCs were resuspended at 20% Ht in PBSG+BSA [PBSG containing 0.5% (w/v) bovine serum albumin (BSA), low endotoxin, ≥98%, pH 7.0, Cod. A1470, Merck Life Sciences S.r.l., Milan, Italy] for a final volume of 20 ml, to which 600 µl of "CD71 MicroBeads" (Miltenyi Biotec, Germany) were added and the mixture was rotated end-over-end for 15 min. Five separation columns of the type LS (Large size, code 130-042-401, Miltenyi Biotec) were inserted in succession, into the permanent magnet of a MidiMACS™ separator (Miltenyi Biotec). Three ml of PBSG+BSA were passed through each column to condition it. Then, 4 ml of RBC suspension containing the "CD71 Microbeads" were loaded on each column to allow $CD71^+$ reticulocytes to bind to the ferromagnetic phase of the column and the unbound cells were collected in the flow-though and saved for the next step. The 4 ml of suspension took ≈15 min to pass thought the column. Three ml PBSG+BSA were then added to wash out the remaining unbound cells and added to the flow-through eluate. The column was extracted from the separator magnet and kept on ice until all the first 5 columns were ready for extraction of the bound reticulocytes. This was performed by adding 1.5 ml of PBSG+BSA to each column and forcing the efflux of the buffer and the reticulocytes, with the aid of the syringe plunger, into a 2.0 ml Eppendorf tube for each of the 5 columns. These samples were spun at 400 x g for 2 min and the reticulocytes carefully resuspended with a small amount of the same buffer in order to pool all the samples in a single 2.0 ml Eppendorf tube that was kept temporarily on ice as a sample of young, immature reticulocytes (**RY**). The flow-through eluate from the first separation, which contained all the RBCs plus residual reticulocytes in a volume larger than the original 20 ml because of addition of the washes, was brought to 20 ml after brief centrifugation and removal of the excess buffer. Cells were carefully resuspended and the suspension was passed again, as 4 ml aliquots, through 5 LS columns inserted sequentially into the magnetic separator. This time, however, the flow rate of the mobile phase through the stationary phase of the column was reduced by attaching a 26G hypodermic needle to the outlet luer-lock of the column, so that the time required for passing 4 ml of RBC suspension at 20% Ht through each column was ≈45 min. These cells were collected and pooled as described before and named **RM**, for mature reticulocytes. Reticulocytes were counted in suitable dilutions, made in duplicate, using a Neubauer haemocytometer, and distributed in aliquots as required for further analyses. The residual reticulocyte-depleted RBC suspension was used for the separation of three populations of RBCs of different age as described below.

**Separation of RBCs into subpopulations of different age**. RBCs of different density were isolated by centrifugation in self-forming Percoll® gradients (Percoll® Plus, GE Healthcare, Milan, Italy) according to Lutz et al.[2], with some modifications. A "20x osmolyte solution" (200 mM $NaH_2PO_4$, 10 mM EDTA, 2.28 M NaCl) was prepared by dissolving the following salts in deionized water and bringing the volume to 200 ml: 5.5196 of $NaH_2PO_4 \bullet H_2O$, 0.744 g of EDTA•$2H_2O$ (di-sodium salt, dihydrated) and 26.649 g of NaCl. The pH of this solution is ≈3.8. A "working suspension" of Percoll® was prepared by weighing 85.4 g of Percoll® Plus in a beaker and adding 5 ml of "20X osmolyte solution". Because the pH of the stock Percoll Plus® suspension is ≈9, after mixing with the osmolyte solution the pH drops to ≈6.8. To this suspension, 3 g of BSA (Code A1470, Merck Life Sciences S.r.l., Milan, Italy) are added and allowed to dissolve completely under mild stirring. The pH is adjusted to 7.4 with 1M NaOH and the volume is brought to 100 ml with deionized water. The final osmolality is ≈315 mosmol/kg $H_2O$. The suspension is stored at 4 °C until use. Two aliquots of the 20% Ht RBC suspension recovered after removing the $CD71^+$ reticulocytes, as described above, and containing each the equivalent of 1.95 ml of packed RBCs were transferred in two polycarbonate centrifuge tubes (Code: 342080, Beckman-Coulter S.r.l., Milan, Italy), which were spun at 1000xg for 5 min at 20 °C in a SS34 rotor of a Sorvall RC-5B centrifuge (Thermo Fisher Scientific, Monza, Italy). The supernatant was carefully aspirated and discarded and to the packed RBCs in each of the two tubes, 11.2 ml of the Percoll® "working suspension" were added. The packed RBCs were carefully and thoroughly mixed with the Percoll®. The final Ht at this point should be < 15%). The tubes were centrifuged for 31 min at 16500 rpm, 20 °C in the same rotor and centrifuge described above. At the end, three fractions were aspirated from the gradients by suction from the top using a capillary tube connected to a peristaltic pump, and collected in three tubes, corresponding to young, middle-age and old erythrocytes (**EY**, **EM** and **EO**). Cells were washed with PBSG and resuspended in the same buffer at ≈20% Ht. The actual Ht was measured using a micro-haematocrit centrifuge and the cells were counted, after suitable dilution, using a Neubauer haemocytometer.

**Sample preparation for Western blotting.** Usually, 50 µl of cell suspensions in PBSG, containing $3x10^7$ reticulocytes or RBCs, were mixed with 450 µl of "Diluted SDS-PAGE sample Buffer" [prepared by mixing 1 volume of 3X SDS-PAGE sample buffer (50 mM Tris/HCl, pH 6.8, 5% SDS (w/v), 35% sucrose (w/v), 5 mM EDTA, 0.01% bromophenol blue, 200 mM dithiotreitol) with 1.7 volumes of 5% SDS (w/v) in MilliQ water] and incubated at 60 °C for 15 min. Aliquots containing 100-150 µl were distributed in separate tubes and stored frozen until analysis. Loading 10 µl of these dissolved samples in SDS-PAGE equals to loading $6x10^5$ cells.

**Preparation of cells for Scanning Electron Microscopy (SEM)**. Aliquots of reticulocyte and RBC suspensions in PBSG, containing $10^7$ cells were packed by centrifugation and the supernatant discarded. Cells were then resuspended with 60 µl of PBS, to which 6 µl of a 2% solution of glutaraldehyde (GA) in PBS were added under gentle swirling. The sample was incubated for 10 min at room temperature with occasional mixing, after which 600 µl of a 6% solution of GA in PBS were added. After 1 h at room temperature, cells were washed repeatedly with deionized water and stored at 4°C. The two GA solutions were prepared immediately before use[3]. To make 1 ml of 2% GA working solution, 50 µl of a commercially available 50%

aqueous solution of GA (Grade I, 50% in $H_2O$, Merck Life Sciences S.r.l., Milan, Italy) that had been stored in frozen aliquots after purchase, were added to 950 µl of PBS. To make 2 ml of the 6% GA working solution 240 µl of the concentrated 50% GA were added to 1.76 ml of PBS. Cells fixed as described above were centrifuged and resuspended in 100 µl of deionized water. Fifty µl of this suspension were transferred to a 500 µl Eppendorf tube to which 50 µl of a 2% aqueous solution of osmium tetroxide (Cod. 75632-5ML, Merck Life Science S.r.l., Milan, Italy) were added. After 30 min at room temperature, cells were sedimented by centrifugation and the supernatant removed. Cells were washed three times with 200 µl of deionized water each time, then 200 µl of 70% ethanol were added. After 30 min cells were sedimented, the supernatant discarded, and 200 µl of 80% ethanol added. Finally, after another 30 min and removal of the ethanol, 200 µl of absolute ethanol were added and the cells left in this condition at 4°C until the next day, when a few µl the suspension of fixed cells in ethanol were transferred to a 5x7 mm silicon chip (Cod. 16007 4" dia. 5 mm x 7 mm diced Silicon Wafer, 187 chips/wafer, Ted Pella Inc, Redding, CA, USA). After evaporation of the solvent, the chip was laid on one side of a bi-adhesive plastic disc coated with conductive material (PELCO Tabs™, Carbon Conductive Tabs, 12mm OD, Code 16084-1 Ted Pella Inc.). The disc was attached to the surface of a standard pin stub mount for SEM (Standard SEM Pin Stub Mount, Ø12.7mm x 8 mm pin height, aluminium, grooved edge, Code 16111 Ted Pella Inc.). The surface of the samples was made electrically conductive by a coating of Pt using a Cressington 208HR Sputter Coater. SEM images were obtained through a FEG-SEM TESCAN Mira3 XMU, Variable Pressure Field Emission Scanning Electron Microscope (TESCAN, Brno, Czech Republic) located at the Arvedi Laboratory, CISRiC, Pavia, Italy. Observations were made, at different magnifications, in secondary electrons mode at 20 kV with an In-Beam SE detector at a working distance of 5 mm.

**Sample preparation for lipid analysis.** Aliquots of reticulocytes and RBCs containing $2x10^7$ cells, were transferred into 1.5 ml Eppendorf tubes and washed 1-2 times with PBSG to remove any trace of BSA. After pelleting by centrifugation, the supernatant was carefully removed and 0.5 ml of ice-cold, HPLC-grade methanol was added. The sample was briefly mixed and then transferred into a 10 ml Pyrex® glass tubes with polyphenolic screw cap (Code 1636/26MP, Italtrade S.r.l., Genoa, Italy). A second 0.5 ml aliquot of methanol was used to rinse the Eppendorf tube and then transferred to the glass tube. Samples were stored at -80°C until analysis.

**Lipid analysis**. The quantitative lipid measurement was done by high resolution mass spectrometry (LTQ-Orbitrap, Thermo Scientific)[4]. Lipids were extracted from cell pellets ($2x10^7$) with an established MTBE (Methyl-tert-butylether) protocol[5]. The Orbitrap Velos Pro hybrid mass spectrometer was operated in Data Dependent Acquisition mode using a HESI II ion source. Full scan profile spectra from m/z 450 to 1050 for positive ion mode and from m/z 400 to 1000 for negative ion mode were acquired in the Orbitrap mass analyser at a resolution of 100k at m/z 400 and < 2 ppm mass accuracy. Samples were measured once in positive polarity and once in negative polarity. For MS/MS experiments, the 10 most abundant ions of the full scan spectrum were sequentially fragmented in the ion trap using He as collision gas (CID, Normalized Collision Energy: 50; Isolation Width: 1.5; Activation Q: 0.2; and Activation Time: 10). Centroided product spectra at a normal scan rate (33 kDa/s) were collected. The custom developed software tool Lipid Data Analyzer was used for data analysis[6,7]. Seven classes of lipids were quantified, together with their subclasses according to the length and number of double bonds of the acyl chains linked to the glycerol or sphingosine moiety: phosphatidylcholine (PC), lysophosphatidylcholine (LPC), sphingomyeline (SM), phosphatidylethanolamine (PE), phosphatidylserine (PS), phosphatidylinositol (PI) and cholesterol (Chol).

**SDS-PAGE and Western blotting**. RBC and reticulocyte proteins were separated by Polyacrylamide Gel Electrophoresis in Sodium DodecylSulfate (SDS-PAGE), following the Laemmli method, in 10% isocratic or 5%-15% gradient polyacrylamide mini-gels (80x60x1.5 mm) in a Mini Protean 3 system (Bio-Rad Laboratories S.r.l., Segrate, Italy). Sample loading is indicated in the Results section or in figure legends.

Proteins separated by the SDS-PAGE were electro-transferred to a PVDF membrane (0.2 μm pores) using a Trans Blot Turbo system (Bio-Rad Laboratories S.r.l., Segrate, Italy) according to manufacturer's instructions. Membranes were blocked with a blocking solution [5% skimmed milk in 20 mM Tris, pH 7.4, 150 mM NaCl, 0.05% Tween 20 (v/v)] and then incubated overnight at 4 °C with the primary antibody directed to the protein of interest.

Antibodies were diluted in a washing buffer [50 mM Tris/HCl pH 7.5, 0.2 M NaCl, 0.5 ml/l Tween 20, 1 g/l polyethylene glycol (PEG) 20000, 1 g/l BSA] that was also used for washing the membranes. After incubation with the appropriate secondary, horseradish-peroxidase-(HRP)-conjugated antibodies, membranes were developed with the chemiluminescence kit Prime Western Blotting Detection Reagent (GE Healthcare, Milan, Italy) and the signal was acquired with a Molecular Imager ChemiDoc XRS+ (Bio-Rad Laboratories S.r.l., Segrate,

Italy). Densitometry of the bands was performed using the software Scion Image (Scion Corporation, USA).

Antibodies used. Mouse monoclonal (BIII-136) anti human Band 3 (B9277, Merck Life Science S.r.l., Milan, Italy). Mouse monoclonal (H68.4) anti human CD71, sc-51829; mouse monoclonal anti human β-spectrin (VD4), sc-53901 (Santa Cruz Biotechnology, Dallas, Texas, USA). HRP-conjugated goat anti-mouse IgGs (170-6516, Bio-Rad Laboratories S.r.l., Segrate, Italy).

#### SUPPLEMENTARY RESULTS

The two populations of $CD71^+$ reticulocytes isolated here in four independent experiments, RY and RM, amounted approximatively to 0.15% and 0.45% of total circulating red cells, respectively. The overall reticulocyte recovery was ≈60%, assuming a conservative, average reticulocyte count of ≈1.0% in normal human subjects[8]. Some reticulocytes (≈10%) could have been lost during leukodepletion[1], or because they presented too low levels of CD71 in the membrane for binding to the microbeads or because, after binding, they detached due to shearing during sample manipulation. A certain percentage of circulating CD71-negative reticulocytes could not in principle be isolated with the method adopted. In the worst case, ≈30% of total reticulocytes (≈0.3% of total blood cells), could have remained mixed with the rest of the RBCs in the sample, and could have been collected together with the population of young RBCs isolated by density gradient centrifugation, being the reticulocytes of low buoyant density[9,10] (see below for a discussion about the possible contamination by reticulocytes of the subpopulation of young RBCs).

**Evaluation of the level of maturity of RY and RM**. The CD71 levels, evaluated by Western blotting, in equal numbers of RY and of RM from four different experiments are shown in **Figure S1A**. The CD71 levels appeared to be strikingly different in the two populations of reticulocytes, so that the chemiluminescence signal was saturated in the RY sample, whereas it was sometimes barely detectable in the RM sample. This obviously prevented quantification by densitometry of the relative CD71 contents in each pair of samples. To overcome the problem a different approach was used. Increasing amounts of RY were loaded in different lanes of a gel together with a single amount of RM in a separate lane of the same gel (**Figure S1B, left**).

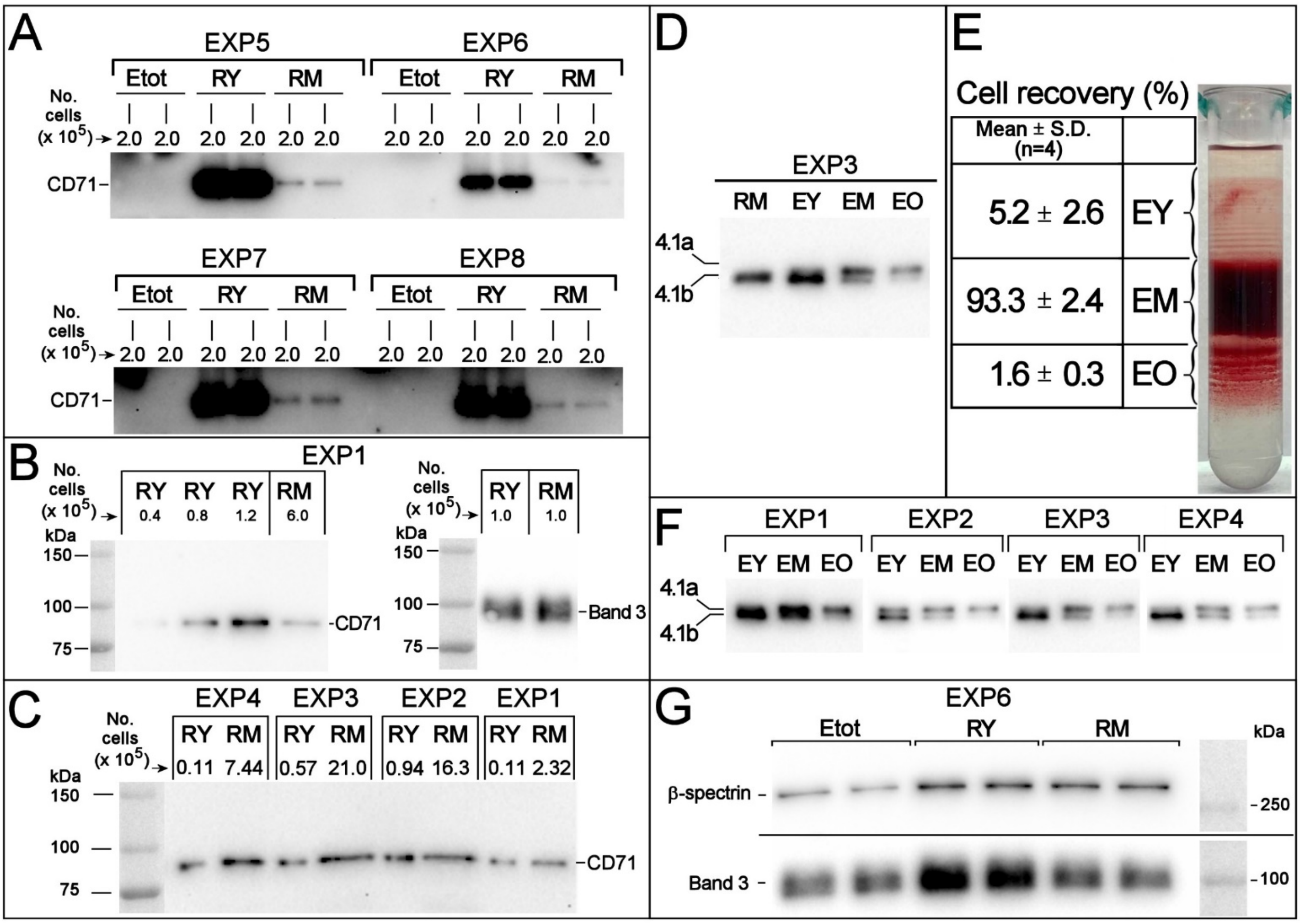


**Figure S1**. **(A)** CD71 levels in RBCs and two populations of reticulocytes. Western blotting with anti CD71 of RY and RM reticulocytes and of total RBCs (Etot) from the corresponding donor in 4 independent experiments. For each sample, 2.0 x $10^5$ cells were loaded. **(B)** Procedure for determining the ratio of CD71 levels between two samples. (Left) CD71 Western blotting of different amounts of RY and one single quantity of RM. (Right) Band 3 Western blotting of identical numbers (1.0x$10^5$) of reticulocytes from the RY and RM populations. Equal amounts of Band 3 confirm the validity of cell count. **(C)** CD71 Western blotting of all the reticulocyte samples obtained in the four experiments. CD71 Western blotting of different amounts of RY and RM from all the experiments. Cell numbers were chosen in order to have the closest possible band intensity for all samples. **(D)** Protein 4.1R Western blotting in reticulocytes and RBCs of different age. The blotting is representative of four independent experiments with identical results. Reticulocytes (RM) and RBCs of different age (EY=young, EM=middle-age, EO=old) were from the same donor. In each lane, 2.0x$10^5$ cells were loaded. See text for details. **(E)** Separation of RBCs according to density in self-forming Percoll® gradients. One tube only is shown of the two that were typically run in each experiment, as representative of four performed experiments. The table gives the average cell recovery in the three RBC subpopulations, evaluated as % Hb (or number of cells) recovered in each fraction with respect to total Hb (or number of cells) loaded in the gradient. **(F)** Western blotting of protein 4.1 in density-separated RBC subpopulations. The relative increase in protein 4.1a/4.1b ratio from young to old RBCs confirms that density separation resulted in a separation of RBCs of different age. Each sample contained 2.0 x $10^5$ cells. **(G)** Band 3 and β-spectrin levels in RY, RM and Etot evaluated by Western blotting. The same number of cells (2x$10^5$) was loaded for each sample, in duplicate. The blot shown is representative of four independent experiments with similar results. See text for details.

After detection of the signal and densitometry of the bands, the RY lane where the CD71 intensity was similar to that of the RM lane was selected. Only when two bands have identical density, the relative CD71 levels in the two samples can be obtained as the ratio between the number of cells loaded for one sample and the number of cells loaded for the other sample. For instance, in the example shown in **Figure S1B**, assuming that the CD71 signal produced by $0.8x10^5$ RY is the same as that of $6.0x10^5$ RM, the ratio of CD71 between RY and RM is 7.5. The same procedure was applied to all four experiments and the results are shown in **Figure S1C**. Because bands were not exactly of the same intensity, an additional correction factor was applied after densitometry of the bands, by linear interpolation between the two closest intensity values of the RY samples, under the assumption of linearity for small intensity differences, to obtain the results shown in **Table S1**.

**Table S1. Ratio between CD71 in RY and in RM**

| **EXPERIMENT** | | **Cell N.** ($x10^5$) | **Band density** (arbitrary units) | **Band density** (%) | **Cell N. after correction** ($x10^5$) | **(CD71 in RY)/ (CD71 in RM)** |
|---|---|---|---|---|---|---|
| **EXP1** | **RY** | 0.114 | 492 | 107.5 | 0.12 | **19** |
| | **RM** | 2.32 | 529 | 100.0 | 2.32 | |
| **EXP2** | **RY** | 0.94 | 1165 | 73.9 | 0.69 | **23** |
| | **RM** | 16.26 | 861 | 100.0 | 16.26 | |
| **EXP3** | **RY** | 0.57 | 712 | 172.8 | 0.99 | **21** |
| | **RM** | 21.00 | 1230 | 100.0 | 21.00 | |
| **EXP4** | **RY** | 0.108 | 627 | 206.5 | 0.22 | **33** |
| | **RM** | 7.44 | 1295 | 100.0 | 7.44 | |

It was also possible to compare the CD71 levels in samples from different donors/experiments, by arbitrarily setting at 100% the CD71 levels of the reticulocyte population from one donor (underscored "100" in **Table S2**) to which the CD levels of the other cell populations could be referred.

**Table S2. Ratio between CD71 in RY and in RM in different donors.**

| **RY** | | | | | | **RM** | | | | | |
|---|---|---|---|---|---|---|---|---|---|---|---|
| EXP. | Cell N. ($x10^5$) | Band density (arbitrary units) | Band density (%) | Cell N. after correction ($x10^5$) | **CD71 levels (% of EXP4)** | EXP. | Cell N. ($x10^5$) | Band density (arbitrary units) | Band density (%) | Cell N. after correction ($x10^5$) | **CD71 levels (% of EXP4)** |
| EXP1 | 0.114 | 492 | 78 | 0.145 | **74** | EXP1 | 2.320 | 529 | 41 | 5.682 | **131** |
| EXP2 | 0.940 | 1165 | 186 | 0.506 | **21** | EXP2 | 16.260 | 861 | 66 | 24.463 | **30** |
| EXP3 | 0.570 | 712 | 114 | 0.502 | **22** | EXP3 | 21.000 | 1230 | 95 | 22.109 | **34** |
| EXP4 | 0.108 | 627 | 100 | 0.108 | **<u>100</u>** | EXP4 | 7.440 | 1295 | 100 | 7.440 | **<u>100</u>** |

**Verification of the purity of the isolated reticulocytes**. To exclude contamination of the reticulocyte samples by mature RBCs, we resorted to analysis by Western blotting of protein 4.1R in reticulocyte samples. Protein 4.1R is synthesized in the erythroid precursors as protein 4.1b, which undergoes a time-dependent, non-enzymatic, post translational deamidation of two asparagine residues that converts it to protein 4.1a[11]. Therefore, protein 4.1a is only present in RBCs, in variable amounts depending on the age of the cell. It is expected that possible contaminant RBCs are from the total population of cells, of intermediate age, where protein 4.1a is present in approximately the same amounts as protein 4.1b. Contamination by RBCs could therefore be revealed by the presence of protein 4.1a in reticulocyte samples even when present in relatively low amounts, starting from a few percent of total cells. As shown in **Figure S1D**, representative of all the performed experiments, no protein 4.1a is visible in the sample of RM reticulocytes, whereas it is clearly detected in the population of young RBCs and progressively increases until becoming the most abundant form in old RBCs (RBCs of different age were obtained from the same donor as described in the next paragraph.)

**Different cell age of density-separated RBC subpopulations.** The average recovery of cells in the three subpopulations of RBCs of different density was: 5.2 2.6 (EY), 93.3 2.4 (EM) and 1.6 0.3 (EO) (n=4) (**Figure S1E**). The effectiveness of the separation of RBCs in subpopulations that differed according to cell age, not only cell density, was confirmed by evaluating the protein 4.1a/4.1b ratio in the different subpopulations by Western blotting. As shown in **Figure S1F**, in all experiments the relative amounts of protein 4.1a increase and those of 4.1b decrease from EY to EO. The EY recovered in the four independent experiments amounted, on average, to ≈5.0% of total cells subjected to density fractionation. If the residual 0.3% (with respect to total blood cells) “missing” reticulocytes (see paragraph “Immuno-magnetic separation of Retics” above) all ended up mixed with EY, 6% of the cells in this fraction would have been reticulocytes, the rest young RBCs. This should not have had a major impact on the characterization of the “true” population of young RBCs (EY), first because of the limited level of contamination, second because of the advanced degree of maturation of CD71-negative reticulocytes.

**Spectrin and Band 3 decrease from RY to EM**. Spectrin and Band 3 are representative of, respectively, peripheral and integral proteins of the RBC membrane. Band 3, together with glycophorin A, the two most abundant integral membrane proteins, appear to be completely retained in the reticulocyte that matures in the bone marrow, at the same time as cells release exosomes enriched in TfR[12]. If the same conservative mechanism that saves Band 3, glycophorin A and membrane-skeletal proteins, at the expense of TfR, were at the basis of the maturation of the circulating reticulocyte, the levels of, i.a., Band 3 and spectrin should be the same when comparing the same number of reticulocytes and young or mature RBCs. **Figure S1G** shows that this is not the case, as both proteins decrease, on a "per cell" basis, from the reticulocyte to the RBC. This strongly supports the concept that marrow reticulocytes and circulating reticulocytes lose membrane area by different mechanisms[1]. Quantification by densitometry reveals that the ratio between Band 3 levels in Retic 15 and RBCs is ≈2.5, pointing to a loss of more than half the total Band 3 content in the process. As the decrease in surface area from circulating reticulocytes to RBCs is expected to be ≈20%[13], this result is clearly absurd, and is related to the non-linear relationship between chemiluminescence emission and protein levels typical of the Western blotting procedure adopted (see above, paragraph "Evaluation of the level of maturity of RY and RM"). However, this artefact proved to be useful here to amplify differences in Band 3 and spectrin levels that would have been otherwise too small to be detected with a linear method.

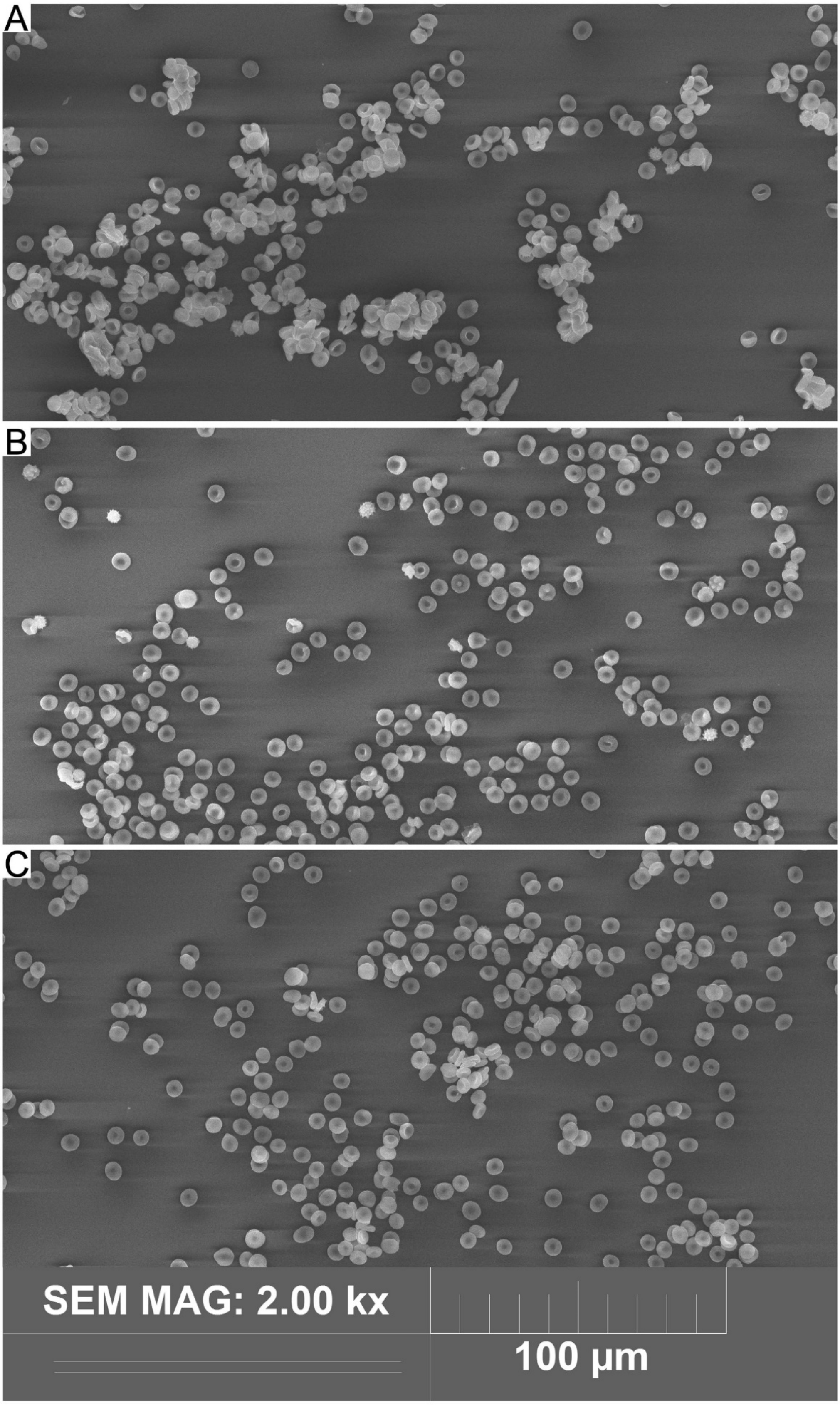


**Figure S2**. **Scanning electron microscopy of reticulocytes and RBCs.** A lower magnification field from reticulocyte samples described in Figure 2. **(A)** RY, **(B)** RM, **(C)**, Etot. Immature reticulocytes did not separate from the CD71-Microbeads as easily as mature reticulocytes when suspended by pipetting, due to the higher amounts of CD71 in their membrane, so they appear aggregated in clumps.

FIGURE S3

| | TOTAL MEMBRANE LIPIDS | | | | |
|---|---|---|---|---|---|
| | RY | RM | EY | EM | EO |
| | Mean ± SD (n=7) | Mean ± SD (n=8) | Mean ± SD (n=8) | Mean ± SD (n=8) | Mean ± SD (n=8) |
| SM | 18.35 ± 0.48 | 18.25 ± 1.21 | 19.13 ± 0.77 | 19.65 ± 1.59 | 18.93 ± 0.41 |
| PC | 23.71 ± 0.09 | 21.84 ± 0.86 | 20.32 ± 1.05 | 21.04 ± 1.27 | 21.67 ± 0.81 |
| LPC | 1.53 ± 0.26 | 1.29 ± 0.09 | 0.96 ± 0.06 | 0.93 ± 0.13 | 0.93 ± 0.10 |
| PE | 9.45 ± 0.84 | 9.47 ± 1.15 | 10.03 ± 2.29 | 10.88 ± 1.53 | 10.05 ± 0.99 |
| PS | 2.87 ± 0.49 | 2.80 ± 0.47 | 2.90 ± 0.53 | 2.63 ± 0.23 | 2.09 ± 0.30 |
| PI | 0.55 ± 0.06 | 0.48 ± 0.07 | 0.32 ± 0.02 | 0.39 ± 0.04 | 0.33 ± 0.03 |
| Chol | 46.68 ± 2.26 | 46.81 ± 2.86 | 47.07 ± 5.75 | 44.48 ± 2.65 | 46.86 ± 2.56 |
| (SM+Chol)/(total lipids) | 0.63 ± 0.02 | 0.64 ± 0.02 | 0.66 ± 0.05 | 0.64 ± 0.03 | 0.65 ± 0.02 |
| (SM+Chol)/glycerophopsholipids | 1.70 ± 0.19 | 1.82 ± 0.18 | 1.94 ± 0.43 | 1.79 ± 0.32 | 1.90 ± 0.20 |

| | p values for the paired t test (all samples n=8 except RY: n=7) | | | | | | | | |
|---|---|---|---|---|---|---|---|---|---|
| | SM/ (total lipids) | PC/ (total lipids) | LPC/ (total lipids) | PE/ (total lipids) | PS/ (total lipids) | PI/ (total lipids) | Chol/ (total lipids) | (SM+Chol)/ (total lipids) | (SM+Chol)/ glycerophospholipids |
| **RY vs. RM** | 0.8424 | 0.0060 | 0.0162 | 0.9287 | 0.9945 | 0.0018 | 0.9567 | 0.5634 | 0.6583 |
| **RY vs. EY** | 0.1190 | 0.0002 | 0.0172 | 0.4872 | 0.8398 | 0.0013 | 0.6522 | 0.0411 | 0.0330 |
| **RY vs. EM** | 0.0342 | 0.0004 | 0.0035 | 0.1017 | 0.6036 | 0.0184 | 0.1858 | 0.3823 | 0.5211 |
| **RY vs. EO** | 0.1740 | 0.0016 | 0.0039 | 0.4768 | 0.0969 | 0.0033 | 0.8670 | 0.0495 | 0.0658 |
| **RM vs. EY** | 0.1940 | 0.0004 | 0.0256 | 0.4846 | 0.6925 | 0.0036 | 0.8127 | 0.2638 | 0.4141 |
| **RM vs. EM** | 0.0703 | 0.0080 | 0.0035 | 0.2681 | 0.5642 | 0.0652 | 0.3123 | 0.8645 | 0.8907 |
| **RM vs. EO** | 0.2530 | 0.6406 | 0.0048 | 0.5054 | 0.0272 | 0.0090 | 0.9671 | 0.5342 | 0.6219 |
| **EY vs. EM** | 0.2460 | 0.0645 | 0.6324 | 0.1906 | 0.1735 | 0.0085 | 0.1017 | 0.1788 | 0.2191 |
| **EY vs. EO** | 0.4504 | 0.0053 | 0.6691 | 0.8837 | 0.0005 | 0.4710 | 0.6240 | 0.3328 | 0.4964 |
| **EM vs. EO** | 0.1263 | 0.0650 | 0.8400 | 0.1610 | 0.0006 | 0.0032 | 0.0610 | 0.2370 | 0.1967 |

**Figure S3**. Tabulated values of membrane lipid content in mol % with respect to total lipids. Mean ± SD of 8 determinations from 4 different donors. The same data are displayed in the graph of next figure **FigureS4** and have been elaborated to generate the regression analysis shown in **Figure 3**.

FIGURE S4

| Paired t test SM/total lipids | p= |
|---|---|
| RY *vs.* RM | 0.8424 |
| RY *vs.* EY | 0.1190 |
| RY *vs.* EM | 0.0342 |
| RY *vs.* EO | 0.1740 |
| RM *vs.* EY | 0.1940 |
| RM *vs.* EM | 0.0703 |
| RM *vs.* EO | 0.2530 |
| EY *vs.* EM | 0.2460 |
| EY *vs.* EO | 0.4504 |
| EM *vs.* EO | 0.1263 |

| Paired t test PC/total lipids | p= |
|---|---|
| RY *vs.* RM | 0.0060 |
| RY *vs.* EY | 0.0002 |
| RY *vs.* EM | 0.0004 |
| RY *vs.* EO | 0.0016 |
| RM *vs.* EY | 0.0004 |
| RM *vs.* EM | 0.0080 |
| RM *vs.* EO | 0.6406 |
| EY *vs.* EM | 0.0645 |
| EY *vs.* EO | 0.0053 |
| EM *vs.* EO | 0.0650 |

| Paired t test LPC/total lipids | p= |
|---|---|
| RY *vs.* RM | 0.0162 |
| RY *vs.* EY | 0.0172 |
| RY *vs.* EM | 0.0035 |
| RY *vs.* EO | 0.0039 |
| RM *vs.* EY | 0.0256 |
| RM *vs.* EM | 0.0035 |
| RM *vs.* EO | 0.0048 |
| EY *vs.* EM | 0.6324 |
| EY *vs.* EO | 0.6691 |
| EM *vs.* EO | 0.8400 |

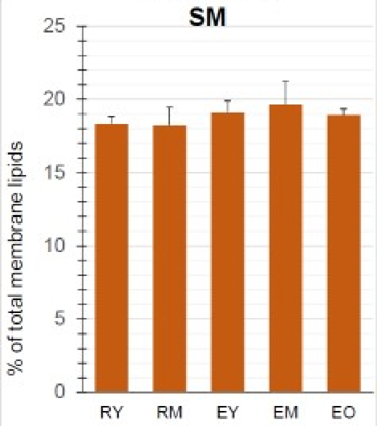


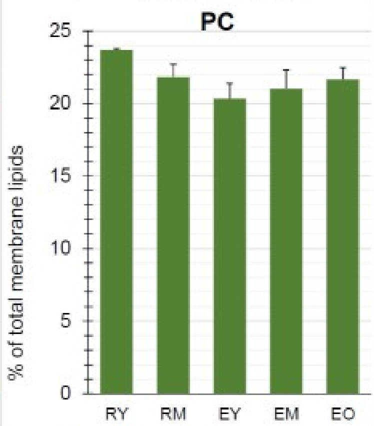


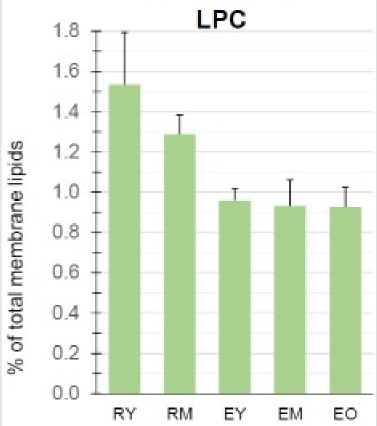


| Paired t test PE/total lipids | p= |
|---|---|
| RY *vs.* RM | 0.9287 |
| RY *vs.* EY | 0.4872 |
| RY *vs.* EM | 0.1017 |
| RY *vs.* EO | 0.4768 |
| RM *vs.* EY | 0.4846 |
| RM *vs.* EM | 0.2681 |
| RM *vs.* EO | 0.5054 |
| EY *vs.* EM | 0.1906 |
| EY *vs.* EO | 0.8837 |
| EM *vs.* EO | 0.1610 |

| Paired t test PS/total lipids | p= |
|---|---|
| RY *vs.* RM | 0.9945 |
| RY *vs.* EY | 0.8398 |
| RY *vs.* EM | 0.6036 |
| RY *vs.* EO | 0.0969 |
| RM *vs.* EY | 0.6925 |
| RM *vs.* EM | 0.5642 |
| RM *vs.* EO | 0.0272 |
| EY *vs.* EM | 0.1735 |
| EY *vs.* EO | 0.0005 |
| EM *vs.* EO | 0.0006 |

| Paired t test PI/total lipids | p= |
|---|---|
| RY *vs.* RM | 0.0018 |
| RY *vs.* EY | 0.0013 |
| RY *vs.* EM | 0.0184 |
| RY *vs.* EO | 0.0033 |
| RM *vs.* EY | 0.0036 |
| RM *vs.* EM | 0.0652 |
| RM *vs.* EO | 0.0090 |
| EY *vs.* EM | 0.0085 |
| EY *vs.* EO | 0.4710 |
| EM *vs.* EO | 0.0032 |

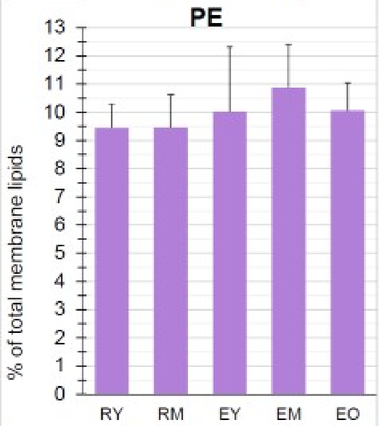


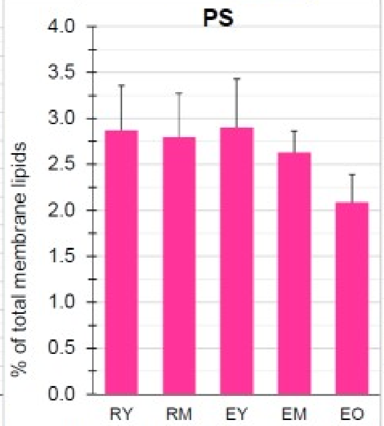


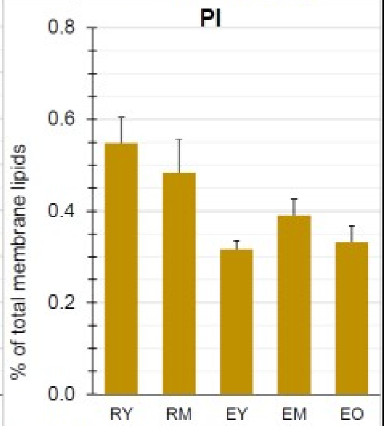


| aired t test Chol/total lipids | p= |
|---|---|
| RY *vs.* RM | 0.9567 |
| RY *vs.* EY | 0.6522 |
| RY *vs.* EM | 0.1858 |
| RY *vs.* EO | 0.8670 |
| RM *vs.* EY | 0.8127 |
| RM *vs.* EM | 0.3123 |
| RM *vs.* EO | 0.9671 |
| EY *vs.* EM | 0.1017 |
| EY *vs.* EO | 0.6240 |
| EM *vs.* EO | 0.0610 |

| Paired t test (SM+Chol)/total lipids | p= |
|---|---|
| RY *vs.* RM | 0.5634 |
| RY *vs.* EY | 0.0411 |
| RY *vs.* EM | 0.3823 |
| RY *vs.* EO | 0.0495 |
| RM *vs.* EY | 0.2638 |
| RM *vs.* EM | 0.8645 |
| RM *vs.* EO | 0.5342 |
| EY *vs.* EM | 0.1788 |
| EY *vs.* EO | 0.3328 |
| EM *vs.* EO | 0.2370 |

| Paired t test (SM+Chol)/glyceroPL | p= |
|---|---|
| RY *vs.* RM | 0.6583 |
| RY *vs.* EY | 0.0330 |
| RY *vs.* EM | 0.5211 |
| RY *vs.* EO | 0.0658 |
| RM *vs.* EY | 0.4141 |
| RM *vs.* EM | 0.8907 |
| RM *vs.* EO | 0.6219 |
| EY *vs.* EM | 0.2191 |
| EY *vs.* EO | 0.4964 |
| EM *vs.* EO | 0.1967 |

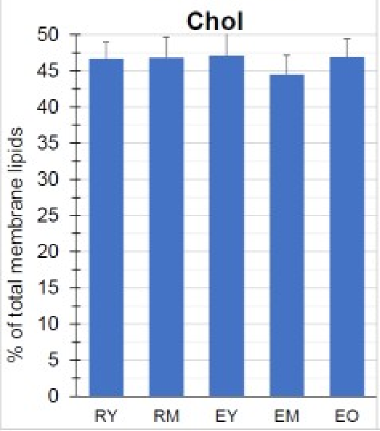


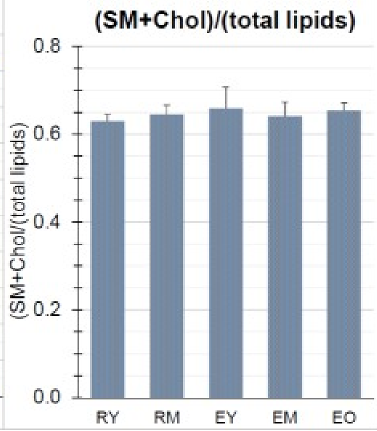


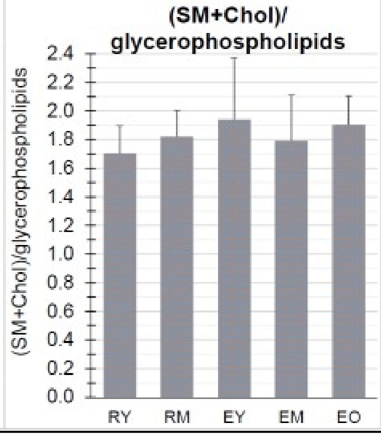

### FIGURE S5

| | PC species (mol% of all PC) | | | | | | |
|---|---|---|---|---|---|---|---|
| | **RY** Mean ± * (n=2) | **RM** Mean ± SD (n=3) | **EY** Mean ± SD (n=4) | **EM** Mean ± SD (n=4) | **EO** Mean ± SD (n=4) | **PLASMA** | **Etot** |
| 30:0 (14:0–16:0) | | | | | | 0.20 | 0.41 |
| 31:0 (15:0–16:0) | | | | | | 0.00 | 0.10 |
| 32:0 (16:0–16:0) | 11.92 ± 0.20 | 12.25 ± 0.72 | 7.63 ± 1.19 | 5.16 ± 0.55 | 4.39 ± 0.47 | 0.81 | 4.70 |
| 32:1 (16:0–16:1) | 1.39 ± 0.14 | 1.30 ± 0.23 | 1.01 ± 0.14 | 0.96 ± 0.15 | 0.98 ± 0.14 | 0.71 | 0.92 |
| 33:0 (16:0–17:0) | | | | | | 0.20 | 0.20 |
| 34:0 (16:0–18:0) | 2.79 ± 0.05 | 2.72 ± 0.14 | 2.25 ± 0.23 | 2.07 ± 0.20 | 1.91 ± 0.21 | 0.20 | 2.15 |
| 34:1 (16:0–18:1ω9) | 39.40 ± 0.85 | 37.26 ± 2.32 | 26.79 ± 2.08 | 22.85 ± 1.01 | 22.57 ± 0.60 | 9.41 | 18.40 |
| 34:1 (16:0–18:1ω7) | | | | | | 2.13 | 2.97 |
| 35:1 (17:0–18:1) | | | | | | 0.40 | 0.41 |
| 35:2 (17:0–18:2) | | | | | | 0.30 | 0.31 |
| 36:0 (18:0–18:0) | | | | | | 0.51 | 0.61 |
| 36:1 (18:0–18:1ω9) | 4.84 ± 0.02 | 4.72 ± 0.65 | 5.78 ± 0.44 | 6.74 ± 0.47 | 6.94 ± 0.52 | 2.13 | 5.93 |
| 36:1 (18:0–18:1ω7) | | | | | | 0.61 | 0.31 |
| 34:2 (16:0–18:2ω6) | 12.79 ± 0.25 | 14.85 ± 1.15 | 22.66 ± 0.85 | 24.16 ± 0.85 | 24.25 ± 1.08 | 29.76 | 27.51 |
| 36:2 (18:1ω9–18:1ω9) | | | | | | 1.32 | 1.84 |
| 36:2 (18:0–18:2ω6) | 7.09 ± 0.03 | 6.91 ± 0.88 | 10.25 ± 1.03 | 13.00 ± 0.89 | 13.44 ± 0.73 | 15.59 | 11.45 |
| 36:3 (16:0–20:3 ) | 4.85 ± 0.15 | 4.91 ± 0.23 | 6.45 ± 0.37 | 7.15 ± 0.40 | 7.46 ± 0.38 | 7.59 | 5.32 |
| 36:3 (18:1ω7–18:2) | | | | | | 1.01 | 0.61 |
| 36:4 (16:0–20:4ω6) | 6.86 ± 0.82 | 7.39 ± 1.16 | 7.73 ± 1.24 | 7.59 ± 1.12 | 7.70 ± 1.15 | 7.79 | 5.21 |
| 36:4 (18:2–18:2 ) | | | | | | 1.92 | 1.02 |
| 37:4 (17:0–20:4) | | | | | | 0.20 | 0.20 |
| 38:3 (18:0–20:3ω6) | 0.54 ± 0.05 | 0.49 ± 0.10 | 0.90 ± 0.21 | 1.23 ± 0.18 | 1.38 ± 0.11 | | |
| 38:4 (18:0–20:4ω6) | 3.47 ± 0.43 | 3.07 ± 0.40 | 3.74 ± 0.53 | 4.01 ± 0.49 | 3.98 ± 0.53 | 6.78 | 3.99 |
| 38:4 (16:0–22:4ω6) | | | | | | 0.91 | 0.51 |
| 38:5 (18:1ω9–20:4) | 1.86 ± 0.04 | 1.80 ± 0.11 | 1.55 ± 0.06 | 1.51 ± 0.15 | 1.52 ± 0.13 | 1.32 | 0.61 |
| 38:5 (18:1ω7–20:4) | | | | | | 0.20 | 0.20 |
| 38:5 (18:0–20:5ω3) | | | | | | 0.30 | 0.31 |
| 38:5 (16:0–22:5ω3) | | | | | | 3.54 | 2.04 |
| 38:6 (16:0–22:6ω3) | | | | | | | |
| 38:6 (18:2–20:4) | 1.87 ± 0.25 | 1.99 ± 0.30 | 2.63 ± 0.82 | 2.77 ± 0.90 | 2.63 ± 0.84 | 0.51 | 0.31 |
| 40:4 (18:0–22:4ω6) | | | | | | 0.40 | 0.41 |
| 40:5 (18:1ω9–22:4ω6) | | | | | | 0.40 | 0.00 |
| 40:6 (18:0–22:5 + 18:0–22:6) | 0.33 ± 0.11 | 0.33 ± 0.12 | 0.62 ± 0.32 | 0.82 ± 0.41 | 0.84 ± 0.42 | 1.52 | 1.02 |
| 40:6 (18:1ω9–22:5 + 18:0–22:6) | | | | | | 1.32 | 0.00 |

*) difference between the mean and each of the two values.

PLASMA and Etot, data from: Myher JJ, Kuksis A, Pind S. Molecular species of glycerophospholipids and sphingomyelins of human erythrocytes: improved method of analysis. Lipids. 1989;24:396-407; and Myher JJ, Kuksis A, Pind S. Molecular species of glycerophospholipids and sphingomyelins of human plasma: comparison to red blood cells. Lipids. 1989;24:408-418.

FIGURE S6

p values for the paired t test (all samples n=4 except RM: n=3 and RY: n=2)

| | 32:0 | 32:1 | 34:0 | 34:1 | 36:1 | 34:2 | 36:2 | 36:3 | 36:4 | 38:3 | 38:4 | 38:5 | 38:6 | 40:6 | (32:0 to 34:1) |
|---|---|---|---|---|---|---|---|---|---|---|---|---|---|---|---|
| RM vs. EY | 0.004 | 0.083 | 0.106 | 0.010 | 0.100 | 0.009 | 0.011 | 0.003 | 0.875 | 0.015 | 0.197 | 0.010 | 0.112 | 0.140 | 0.007 |
| RM vs. EM | 0.000 | 0.034 | 0.054 | 0.005 | 0.052 | 0.004 | 0.001 | 0.001 | 0.180 | 0.006 | 0.144 | 0.038 | 0.111 | 0.090 | 0.002 |
| RM vs. EO | 0.001 | 0.038 | 0.020 | 0.006 | 0.033 | 0.005 | 0.001 | 0.005 | 0.905 | 0.002 | 0.167 | 0.009 | 0.081 | 0.072 | 0.002 |
| EY vs. EM | 0.006 | 0.049 | 0.009 | 0.007 | 0.022 | 0.082 | 0.001 | 0.006 | 0.590 | 0.001 | 0.059 | 0.396 | 0.051 | 0.030 | 0.005 |
| EY vs. EO | 0.004 | 0.160 | 0.007 | 0.013 | 0.001 | 0.055 | 0.002 | 0.011 | 0.876 | 0.003 | 0.178 | 0.347 | 0.992 | 0.037 | 0.007 |
| EM vs. EO | 0.001 | 0.199 | 0.054 | 0.291 | 0.521 | 0.709 | 0.039 | 0.050 | 0.486 | 0.032 | 0.836 | 0.838 | 0.273 | 0.391 | 0.029 |

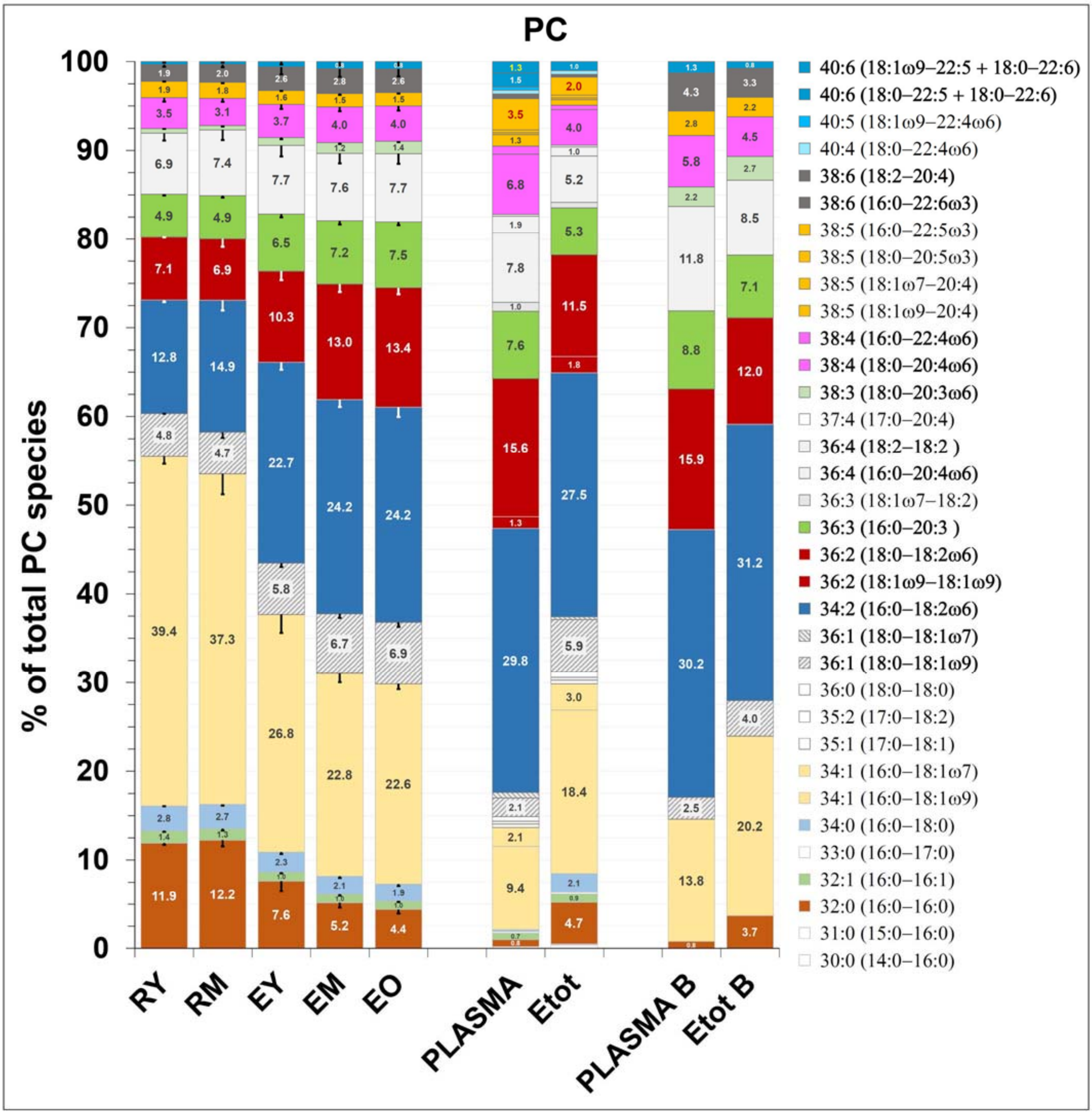


**Figure S6**. **PLASMA** and **Etot**, data from: Myher JJ, Kuksis A, Pind S. Molecular species of glycerophospholipids and sphingomyelins of human erythrocytes: improved method of analysis. Lipids. 1989;24:396-407; Myher JJ, Kuksis A, Pind S. Molecular species of glycerophospholipids and sphingomyelins of human plasma: comparison to red blood cells. Lipids. 1989;24:408-418.

**PLASMA B** and **Etot B**, data from: Dushianthan A, Cusack R, Koster G, Grocott MPW, Postle AD. Insight into erythrocyte phospholipid molecular flux in healthy humans and in patients with acute respiratory distress syndrome. PLoS One. 2019;14:e0221595.

Error bars represent the standard deviation: EY, EM, EO, n=4; RM, n=3; RY, n=2. For RY, where only two samples were available, the error bars represent the difference between the mean and the individual values.

| | Diacyl PE species (mol% of all diacyl PE) | | | | | | |
|---|---|---|---|---|---|---|---|
| | RY<br>Mean ± * (n=2) | RM<br>Mean ± SD (n=3) | EY<br>Mean ± SD (n=4) | EM<br>Mean ± SD (n=4) | EO<br>Mean ± SD (n=4) | PLASMA | Etot |
| 26:0 (14:0-16:0) | 0.81 ± 0.02 | 1.08 ± 0.13 | 1.10 ± 0.22 | 0.96 ± 0.20 | 1.18 ± 0.24 | | |
| 31:0 (15:0-16:0) | | | | | | | |
| 32:0 (16:0-16:0) | | | | | | 0.32 | 0.62 |
| 32:1 (16:0-16:1) | 2.94 ± 1.31 | 2.29 ± 2.68 | 2.13 ± 1.25 | 0.41 ± 0.19 | 0.47 ± 0.09 | 0.32 | 0.41 |
| 33:0 (16:0-17:0) | | | | | | | |
| 34:0 (16:0-18:0) | | | | | | 0.21 | 0.31 |
| 34:1 (16:0-18:1w9) | 13.30 ± 1.08 | 13.77 ± 1.36 | 17.76 ± 2.81 | 21.95 ± 2.02 | 23.24 ± 2.11 | 2.99 | 15.66 |
| 34:1 (16:0-18:1w7) | | | | | | 0.75 | 1.04 |
| 34:2 (16:0-18:2w6) | 6.68 ± 2.00 | 6.00 ± 3.85 | 5.94 ± 0.91 | 5.77 ± 1.08 | 6.79 ± 1.36 | 8.24 | 6.43 |
| 35:1 (17:0-18:1) | | | | | | 0.11 | 0.31 |
| 35:2 (17:0-18:2) | | | | | | 0.32 | 0.31 |
| 36:0 (18:0-18:0) | | | | | | 0.32 | 0.10 |
| 36:1 (18:0-18:1w9) | 3.51 ± 0.04 | 4.04 ± 0.41 | 4.73 ± 0.47 | 5.68 ± 0.22 | 5.82 ± 0.36 | 2.78 | 4.46 |
| 36:1 (18:0-18:1w7) | | | | | | 0.32 | 0.21 |
| 36:2 (18:1w9-18:1w9) | 6.13 ± 0.19 | 6.16 ± 0.66 | 7.36 ± 1.13 | 9.82 ± 0.14 | 10.94 ± 0.24 | 2.03 | 4.77 |
| 36:2 (18:1w9-18:1w7) | | | | | | | 0.52 |
| 36:2 (18:0-18:2) | | | | | | 14.76 | 2.18 |
| 36:3 (16:0-20:3 ) | 3.65 ± 0.29 | 3.77 ± 0.35 | 4.87 ± 1.08 | 6.93 ± 0.70 | 8.45 ± 1.09 | 6.31 | 3.63 |
| 36:3 (18:1w9-18:2) | | | | | | | |
| 36:3 (18:1w7-18:2) | | | | | | 0.53 | |
| 36:4 (16:0-20:4w6) | 11.72 ± 0.32 | 12.53 ± 1.30 | 13.72 ± 0.75 | 13.15 ± 0.73 | 12.39 ± 0.95 | 8.02 | 14.00 |
| 36:4 (18:2-18:2 ) | | | | | | 0.75 | 0.93 |
| 36:5 (16:0-20:5) | | | | | | | |
| 37:4 (17:0-20:4) | | | | | | 0.32 | 0.41 |
| 38:4 (18:0-20:4) | 23.99 ± 1.96 | 22.60 ± 1.76 | 18.40 ± 2.47 | 13.99 ± 1.78 | 11.80 ± 1.83 | 24.28 | 13.07 |
| 38:3 (18:0-20:3w6) | | | | | | | |
| 38:4 (16:0-22:4w6) | | | | | | 2.89 | 5.39 |
| 38:4 (18:1-20:3 + 18:1t-20:4) | | | | | | | |
| 38:5 (18:1w9-20:4) | 12.94 ± 1.46 | 13.92 ± 2.95 | 12.30 ± 1.36 | 11.21 ± 1.38 | 10.00 ± 1.44 | 5.88 | 8.71 |
| 38:5 (18:1w7-20:4) | | | | | | 0.75 | 0.93 |
| 38:5 (18:0-20:5w3) | | | | | | 1.28 | 0.62 |
| 38:5 (16:0-22:5w3) | | | | | | 8.34 | 4.88 |
| 38:6 (16:0-22:6w3) | | | | | | | |
| 38:6 (18:2-20:4) | 5.86 ± 1.60 | 5.54 ± 1.58 | 4.80 ± 1.26 | 4.55 ± 1.25 | 4.31 ± 1.19 | 1.18 | 1.87 |
| 40:4 (18:0-22:4w6) | 1.49 ± 0.46 | 1.46 ± 0.37 | 1.35 ± 0.27 | 1.07 ± 0.23 | 0.88 ± 0.21 | 1.18 | 1.97 |
| 40:5 (18:0-22:5w6) | 2.20 ± 0.06 | 2.25 ± 0.35 | 1.83 ± 0.19 | 1.46 ± 0.28 | 1.11 ± 0.23 | | 0.73 |
| 40:5 (18:1w9-22:4w6) | | | | | | | 1.35 |
| 40:6 (18:0-22:5 + 18:0-22:6) | 2.20 ± 0.45 | 1.98 ± 0.48 | 1.67 ± 0.44 | 1.23 ± 0.30 | 0.87 ± 0.17 | 4.81 | 2.07 |
| 40:6 () | | | | | | | 0.21 |
| 40:6 (18:2-22:4) | | | | | | | 0.21 |
| 40:6 (18:1w9-22:5 + 18:0-22:6) | | | | | | | 1.66 |
| 40:7 () | 2.55 ± 0.86 | 2.60 ± 0.87 | 2.04 ± 0.82 | 1.82 ± 0.65 | 1.74 ± 0.64 | | |

**Figure S7. PLASMA** and **Etot**, data from: Myher JJ, Kuksis A, Pind S. Molecular species of glycerophospholipids and sphingomyelins of human erythrocytes: improved method of analysis. Lipids. 1989;24:396-407; Myher JJ, Kuksis A, Pind S. Molecular species of glycerophospholipids and sphingomyelins of human plasma: comparison to red blood cells. Lipids. 1989;24:408-418.
Error bars represent the standard deviation: EY, EM, EO, n=4; RM, n=3; RY, n=2. For RY, where only two samples were available, the error bars represent the difference between the mean and the individual values.

### FIGURE S8

**SM species (mol% of all SM)**

| | RY | RM | EY | EM | EO | | | | | | |
|---|---|---|---|---|---|---|---|---|---|---|---|
| | Mean ± * (n=2) | Mean ± SD (n=3) | Mean ± SD (n=4) | Mean ± SD (n=4) | Mean ± SD (n=4) | Chylos (a) | VLDL (a) | VLDL( b) | LDL (b) | HDL3 (b) | Etot |
| 32:0 () | | | | | | 0.05 | 0.04 | 0.04 | 0.04 | 0.07 | 0.03 |
| d32:1 (d16:1-16:0) | 1.84 ± 0.02 | 1.86 ± 0.05 | 1.96 ± 0.06 | 2.02 ± 0.08 | 2.09 ± 0.05 | 1.36 | 1.29 | 0.98 | 1.08 | 0.17 | 0.92 |
| d32:2 (d16:1-16:1) | | | | | | 0.06 | 0.05 | 0.02 | 0.01 | 0.00 | 0.00 |
| d32:1 (d17:1-15:0) | | | | | | 0.01 | 0.01 | 0.00 | 0.01 | 0.03 | 0.01 |
| d32:1 (d18:1-14:0) | | | | | | 0.74 | 0.69 | 0.50 | 0.61 | 1.00 | 0.36 |
| d32:2 (d18:2-14:0) | | | | | | 0.15 | 0.11 | 0.06 | 0.06 | 0.31 | 0.05 |
| 33:0 () | | | | | | 0.09 | 0.00 | 0.01 | 0.00 | 0.01 | 0.00 |
| d33:1 (d16:1-17:0) | | | | | | 0.08 | 0.07 | 0.02 | 0.01 | 0.03 | 0.04 |
| d33:1 (d17:1-16:0) | 1.30 ± 0.07 | 1.39 ± 0.11 | 1.51 ± 0.16 | 1.55 ± 0.17 | 1.59 ± 0.17 | 1.44 | 1.40 | 0.95 | 0.50 | 0.40 | 0.90 |
| d33:2 (d17:1-16:1) | | | | | | 0.03 | 0.03 | 0.01 | 0.00 | 0.01 | 0.01 |
| d33:1 (d18:1-15:0) | | | | | | 0.08 | 0.08 | 0.00 | 0.03 | 0.05 | 0.13 |
| d33:2 (d18:2-15:0) | | | | | | 0.05 | 0.03 | 0.00 | 0.00 | 0.02 | 0.01 |
| 34:0 () | 0.97 ± 0.09 | 1.08 ± 0.18 | 1.19 ± 0.16 | 1.28 ± 0.20 | 1.29 ± 0.19 | 0.67 | 0.68 | 1.17 | 1.06 | 0.38 | 0.73 |
| d34:1 (d16:1-18:0) | | | | | | 1.88 | 2.03 | 1.81 | 1.34 | 1.54 | 1.40 |
| d34:1 (d16:1-18:1) | | | | | | 0.28 | 0.27 | 0.19 | 0.19 | 0.22 | 0.16 |
| d34:1 (d17:1-17:0) | | | | | | 0.01 | 0.05 | 0.06 | 0.00 | 0.03 | 0.04 |
| d34:1 (d18:1-16:0) | 20.84 ± 0.95 | 21.42 ± 1.72 | 22.81 ± 1.69 | 23.19 ± 1.27 | 23.71 ± 1.90 | 30.36 | 30.12 | 30.46 | 28.77 | 17.29 | 25.87 |
| d34:2 (d18:1-16:1) | | | | | | 0.39 | 0.39 | 0.27 | 0.31 | 0.30 | 0.21 |
| d34:2 (d18:2-16:0) | 1.65 ± 0.02 | 1.65 ± 0.02 | 1.79 ± 0.12 | 1.90 ± 0.15 | 1.95 ± 0.12 | 4.04 | 3.81 | 3.25 | 2.24 | 3.33 | 1.54 |
| d34:3 (d18:2-16:1) | | | | | | 0.00 | 0.00 | 0.00 | 0.00 | 0.00 | 0.00 |
| 35:0 () | | | | | | 0.17 | 0.03 | 0.00 | 0.02 | 0.16 | 0.17 |
| d35:1 (d16:1-19:0) | | | | | | 0.24 | 0.15 | 0.00 | 0.00 | 0.10 | 0.06 |
| d35:1 (d17:1-18:0) | | | | | | 0.66 | 0.54 | 0.73 | 0.54 | 0.43 | 0.30 |
| d35:2 (d17:1-18:1) | | | | | | 0.12 | 0.05 | 0.07 | 0.03 | 0.01 | 0.01 |
| d35:1 (d18:1-17:0) | | | | | | 0.93 | 0.61 | 0.75 | 0.84 | 0.24 | 0.51 |
| d35:2 (d18:2-17:0) | | | | | | 0.19 | 0.16 | 0.06 | 0.08 | 0.20 | 0.05 |
| 36:0 () | | | | | | 0.22 | 0.27 | 0.27 | 0.20 | 0.17 | 0.08 |
| d36:1 (d16:1-20:0) | | | | | | 0.94 | 1.11 | 1.17 | 1.01 | 0.90 | 0.67 |
| d36:1 (d18:1-18:0) | 1.69 ± 0.0002 | 1.81 ± 0.07 | 2.74 ± 0.24 | 3.63 ± 0.17 | 3.98 ± 0.21 | 5.68 | 6.27 | 4.67 | 4.20 | 2.97 | 2.23 |
| d36:2 (d16:1-20:1) | | | | | | 0.12 | 0.10 | 0.08 | 0.11 | 0.03 | 0.05 |
| d36:2 (d18:1-18:1) | | | | | | 0.64 | 0.74 | 0.69 | 0.54 | 0.67 | 0.25 |
| d36:2 (d18:2-18:0) | 0.65 ± 0.01 | 0.75 ± 0.08 | 1.15 ± 0.13 | 1.30 ± 0.09 | 1.35 ± 0.07 | 2.46 | 2.51 | 1.81 | 1.34 | 1.94 | 0.59 |
| d36:3 (d18:2-18:1) | | | | | | 0.33 | 0.34 | 0.21 | 0.16 | 0.31 | 0.05 |
| 37:0 () | | | | | | 0.03 | 0.04 | 0.06 | 0.03 | 0.03 | 0.00 |
| 37:1 () | | | | | | 0.76 | 0.89 | 0.74 | 0.68 | 0.47 | 0.15 |
| 37:2 () | | | | | | 0.20 | 0.19 | 0.00 | 0.00 | 0.00 | 0.00 |
| 38:0 () | | | | | | 0.06 | 0.07 | 0.13 | 0.12 | 0.07 | 0.02 |
| d38:1 (d16:1-22:0) | 1.31 ± 0.04 | 1.02 ± 0.17 | 1.02 ± 0.10 | 1.16 ± 0.12 | 1.26 ± 0.16 | 0.90 | 1.04 | 1.19 | 1.73 | 2.01 | 0.13 |
| d38:2 (d16:1-22:1) | | | | | | 0.08 | 0.07 | 0.13 | 0.17 | 0.08 | 0.00 |
| d38:1 (d17:1-21:0) | | | | | | 0.05 | 0.04 | 0.10 | 0.04 | 0.06 | 0.76 |
| d38:1 (d18:1-20:0) | | | | | | 1.83 | 2.07 | 1.43 | 2.23 | 2.28 | 0.22 |
| d38:2 (d18:1-20:1) | | | | | | 0.12 | 0.08 | 0.13 | 0.16 | 0.13 | 0.02 |
| d38:2 (d18:2-20:0) | | | | | | 0.87 | 0.96 | 0.76 | 0.80 | 1.72 | 0.08 |
| d38:3 (d18:2-20:1) | | | | | | 0.31 | 0.13 | 0.05 | 0.11 | 0.14 | 0.01 |
| 39:0 () | | | | | | 0.02 | 0.01 | 0.02 | 0.01 | 0.07 | 0.00 |
| d39:1 (d16:1-23:0) | | | | | | 0.34 | 0.35 | 0.51 | 0.71 | 0.60 | 0.06 |
| d39:2 (d16:1-23:1) | | | | | | 0.07 | 0.01 | 0.04 | 0.05 | 0.12 | 0.00 |
| d39:1 (d17:1-22:0) | | | | | | 0.48 | 0.39 | 0.51 | 0.62 | 0.62 | 0.12 |
| d39:2 (d17:1-22:1) | | | | | | 0.05 | 0.05 | 0.08 | 0.09 | 0.09 | 0.02 |
| d39:1 (d18:1-21:0) | | | | | | 0.26 | 0.23 | 0.15 | 0.25 | 0.24 | 0.07 |
| d39:2 (d18:2-21:0) | | | | | | 0.19 | 0.14 | 0.09 | 0.09 | 0.34 | 0.03 |
| 40:0 () | | | | | | 0.06 | 0.04 | 0.16 | 0.17 | 0.04 | 0.02 |
| d40:1 (d16:1-24:0) | 8.45 ± 0.10 | 8.03 ± 0.72 | 7.85 ± 0.39 | 7.58 ± 0.63 | 7.27 ± 0.78 | 0.17 | 0.26 | 0.32 | 0.73 | 1.13 | 0.92 |
| d40:1 (d17:1-23:0) | | | | | | 0.40 | 0.43 | 0.24 | 0.34 | 0.44 | 0.01 |
| d40:1 (d18:1-22:0) | | | | | | 5.72 | 5.10 | 5.48 | 8.12 | 8.83 | 4.73 |
| d40:2 (d16:1-24:1) | 3.45 ± 0.13 | 3.22 ± 0.27 | 2.87 ± 0.29 | 2.85 ± 0.21 | 2.94 ± 0.29 | 1.25 | 1.32 | 1.50 | 1.95 | 2.63 | 1.11 |
| d40:2 (d17:1-23:1) | | | | | | 0.07 | 0.10 | 0.02 | 0.07 | 0.05 | 0.00 |
| d40:2 (d18:1-22:1) | | | | | | 0.11 | 0.16 | 0.64 | 0.66 | 0.42 | 0.19 |
| d40:2 (d18:2-22:0) | | | | | | 1.90 | 1.83 | 2.36 | 2.26 | 4.71 | 0.78 |
| d40:3 (d18:2-22:1) | | | | | | 0.46 | 0.46 | 0.39 | 0.63 | 0.85 | 0.19 |
| 41:0 () | | | | | | 0.01 | 0.01 | 0.00 | 0.01 | 0.00 | 0.09 |
| d41:1 (d17:1-24:0) | 2.00 ± 0.21 | 1.89 ± 0.16 | 1.88 ± 0.15 | 1.85 ± 0.17 | 1.79 ± 0.19 | 0.36 | 0.34 | 0.44 | 0.49 | 0.62 | 0.42 |
| d41:1 (d18:1-23:0) | | | | | | 2.10 | 2.05 | 2.63 | 3.05 | 2.81 | 0.24 |
| d41:2 (d17:1-24:1) | 1.23 ± 0.14 | 1.19 ± 0.08 | 1.07 ± 0.07 | 1.10 ± 0.07 | 1.13 ± 0.08 | 0.55 | 0.55 | 0.69 | 0.69 | 1.00 | 0.41 |
| d41:2 (d18:1-23:1) | | | | | | 0.07 | 0.22 | 0.31 | 0.27 | 0.38 | 0.08 |
| d41:2 (d18:2-23:0) | | | | | | 0.77 | 0.76 | 0.89 | 0.82 | 1.46 | 0.39 |
| d41:3 (d17:1-24:2) | | | | | | 0.05 | 0.06 | 0.05 | 0.08 | 0.08 | 0.03 |
| d41:3 (d18:2-23:1 ) | | | | | | 0.22 | 0.16 | 0.08 | 0.14 | 0.24 | 0.00 |
| 42:0 () | | | | | | | 0.00 | 0.04 | 0.00 | 0.01 | 0.04 |
| d42:1 (d18:1-24:0) | 18.00 ± 0.94 | 17.88 ± 0.19 | 18.38 ± 1.38 | 18.14 ± 1.36 | 17.80 ± 1.60 | 5.11 | 4.85 | 5.05 | 6.36 | 5.23 | 19.70 |
| d42:2 (d18:1-24:1) | 27.22 ± 0.70 | 27.76 ± 1.22 | 25.75 ± 1.50 | 24.88 ± 1.54 | 24.36 ± 1.19 | 12.69 | 13.00 | 12.95 | 12.18 | 13.41 | 20.75 |
| d42:2 (d18:2-24:0) | | | | | | 2.24 | 2.15 | 3.20 | 1.61 | 4.59 | 5.28 |
| d42:3 (d18:1-24:2) | 7.04 ± 0.25 | 6.72 ± 0.41 | 5.74 ± 0.50 | 5.33 ± 0.25 | 5.37 ± 0.50 | 0.37 | 0.56 | 0.95 | 1.00 | 2.04 | 2.31 |
| d42:3 (d18:2-24:1) | | | | | | 3.35 | 4.11 | 3.39 | 3.26 | 4.80 | 0.00 |
| d42:4 (d18:2-24:2) | | | | | | 0.88 | 0.65 | 0.33 | 0.40 | 0.00 | 0.39 |
| 43:0 () | | | | | | 0.00 | 0.00 | 1.50 | 1.51 | 1.80 | 0.80 |
| 44:0 () | | | | | | 0.00 | 0.00 | 0.00 | 0.00 | 0.00 | 2.01 |
| d44:1 (d44:1) | 1.08 ± 0.14 | 1.05 ± 0.13 | 1.13 ± 0.16 | 1.09 ± 0.15 | 1.02 ± 0.16 | | | | | | |
| d44:2 (d44:2) | 1.29 ± 0.19 | 1.28 ± 0.17 | 1.14 ± 0.20 | 1.17 ± 0.16 | 1.11 ± 0.17 | | | | | | |

*) difference between the mean and each of the two values.

**p values for the paired t test (all samples n=4 except RM: n=3 and RY: n=2)**

| | d32:1 | d33:1 | d34:0 | d34:1 | d34:2 | d36:1 | d36:2 | d38:1 | d40:1 | d40:2 | d41:1 | d41:2 | d42:1 | d42:2 | d42:3 | d44:1 | d44:2 | (d32:1 to d36:2) |
|---|---|---|---|---|---|---|---|---|---|---|---|---|---|---|---|---|---|---|
| RM *vs.* EY | 0.138 | 0.140 | 0.017 | 0.392 | 0.028 | 0.009 | 0.005 | 0.649 | 0.313 | 0.100 | 0.156 | 0.077 | 0.803 | 0.263 | 0.001 | 0.232 | 0.223 | 0.139 |
| RM *vs.* EM | 0.098 | 0.089 | 0.041 | 0.220 | 0.060 | 0.000 | 0.001 | 0.272 | 0.066 | 0.042 | 0.151 | 0.175 | 0.951 | 0.161 | 0.036 | 0.507 | 0.449 | 0.054 |
| RM *vs.* EO | 0.007 | 0.065 | 0.048 | 0.273 | 0.012 | 0.001 | 0.001 | 0.137 | 0.020 | 0.160 | 0.133 | 0.160 | 0.922 | 0.015 | 0.059 | 0.971 | 0.111 | 0.058 |
| EY *vs.* EM | 0.243 | 0.005 | 0.037 | 0.183 | 0.020 | 0.001 | 0.008 | 0.001 | 0.142 | 0.715 | 0.194 | 0.113 | 0.193 | 0.021 | 0.140 | 0.033 | 0.449 | 0.007 |
| EY *vs.* EO | 0.044 | 0.038 | 0.097 | 0.029 | 0.007 | 0.000 | 0.008 | 0.009 | 0.060 | 0.303 | 0.066 | 0.016 | 0.461 | 0.042 | 0.203 | 0.008 | 0.285 | 0.001 |
| EM *vs.* EO | 0.081 | 0.117 | 0.863 | 0.242 | 0.354 | 0.003 | 0.030 | 0.036 | 0.058 | 0.127 | 0.035 | 0.172 | 0.597 | 0.361 | 0.807 | 0.063 | 0.004 | 0.024 |

Sample **Etot**, data from: Myher JJ, Kuksis A, Pind S. Molecular species of glycerophospholipids and sphingomyelins of human erythrocytes: improved method of analysis. Lipids. 1989;24:396-407. doi: 10.1007/BF02535147

Samples of lipoproteins, data from: Myher JJ, Kuksis A, Pind S. Molecular species of glycerophospholipids and sphingomyelins of human plasma: comparison to red blood cells. Lipids. 1989;24:408-418. doi: 10.1007/BF02535148. (a) Postprandial plasma of a normolipemic subject (b) Fasting plasma from three different normolipemic subjects.

SM is ≈18% of total PL in plasma and ≈25% of total PL in RBCs.

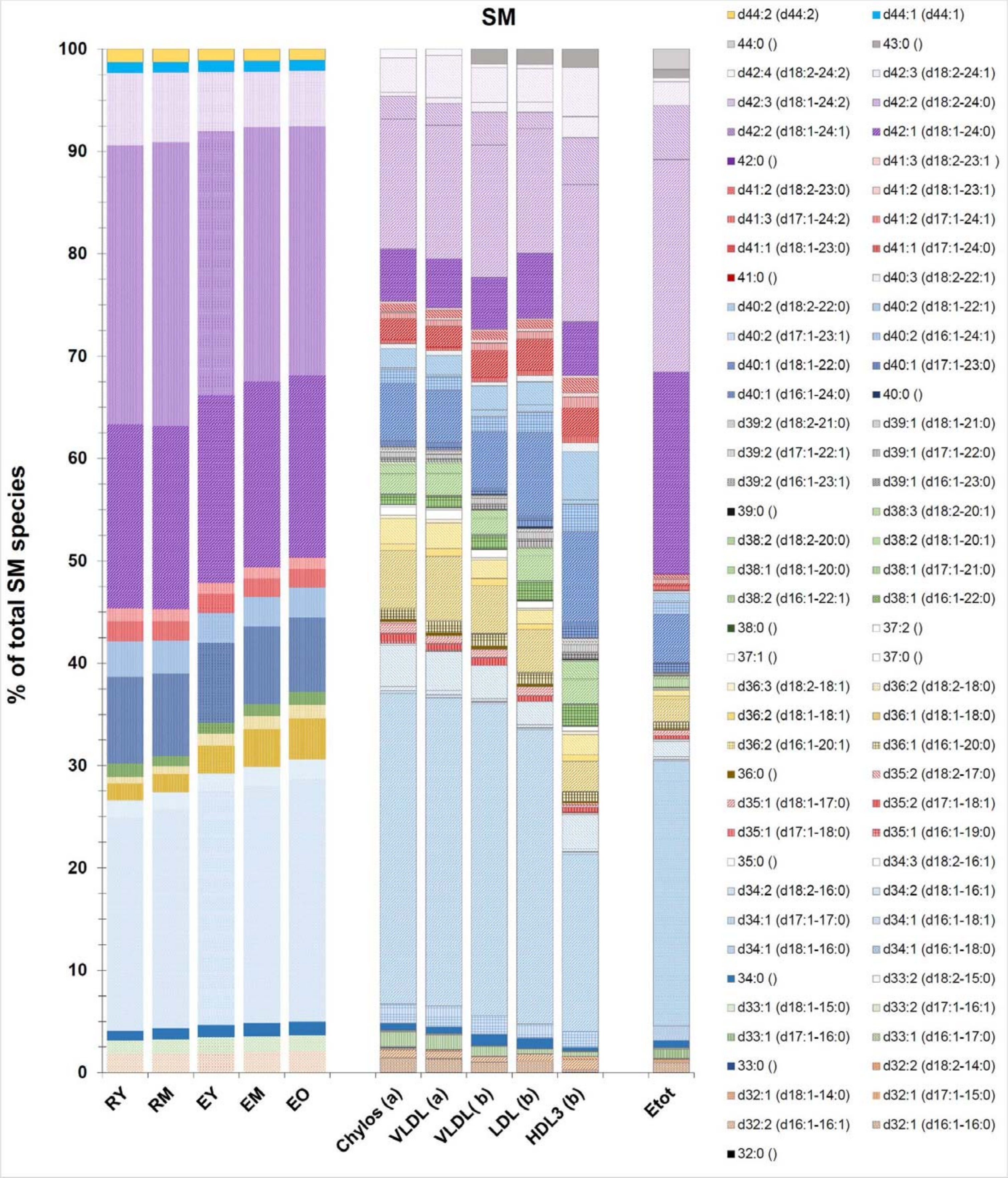


Figure S9. Sample **Etot**, data from: Myher JJ, Kuksis A, Pind S. Molecular species of glycerophospholipids and sphingomyelins of human erythrocytes: improved method of analysis. Lipids. 1989;24:396-407. doi: 10.1007/BF02535147. Samples of lipoproteins, data from: Myher JJ, Kuksis A, Pind S. Molecular species of glycerophospholipids and sphingomyelins of human plasma: comparison to red blood cells. Lipids. 1989;24:408-418. doi: 10.1007/BF02535148. (a) Postprandial plasma of a normolipemic subject (b) Fasting plasma from three different normolipemic subjects. SM is ≈18% of total phospholipids in plasma and ≈25% of total phospholipids in RBCs.

### FIGURE S10

| paired t test<br>(SM d32:1 to SM d36:2) vs. (SM d38:1 to SM d44:2) | | paired t test<br>SATURATED *vs.* UNSATURATED | |
|---|---|---|---|
| RM *vs.* EY | p= 0.1385 | RM *vs.* EY | p= 0.1610 |
| RM *vs.* EM | p= 0.0543 | RM *vs.* EM | p= 0.0731 |
| RM *vs.* EO | p= 0.0577 | RM *vs.* EO | p= 0.0184 |
| EY *vs.* EM | p= 0.0068 | EY *vs.* EM | p= 0.0657 |
| EY *vs.* EO | p= 0.0013 | EY *vs.* EO | p= 0.0952 |
| EM vs. EO | p= 0.0236 | EM vs. EO | p= 0.4355 |

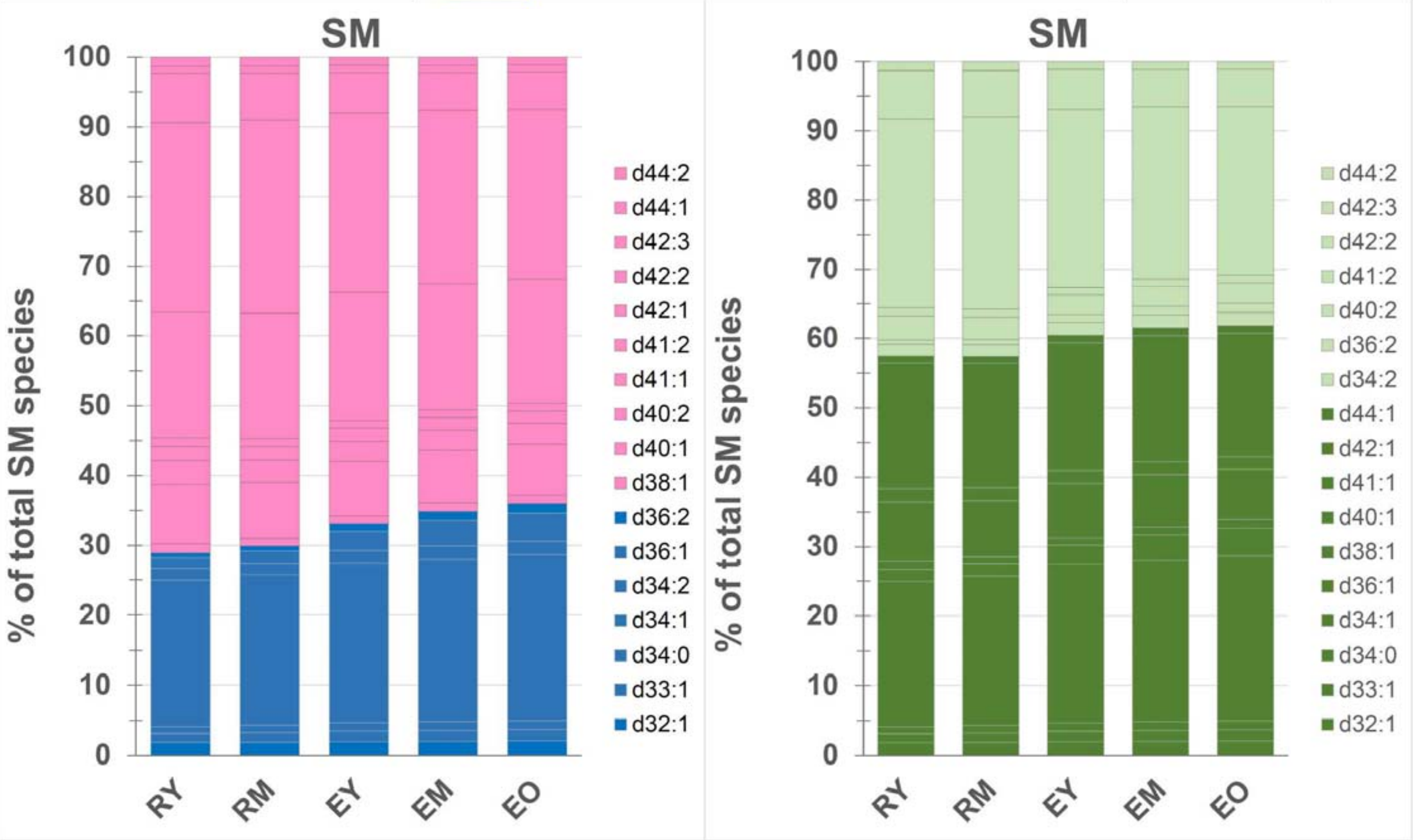


**Figure S10. Left**: the histogram shows in blue the relative abundance with respect of total SM of those species where the amidated acyl chain is shorter than 20 carbon atoms (from d32:1 to D36:1 included) and in pink the sum of the remaining SM subspecies. A general trend to increase is clearly visible for the "short" species during maturation of reticulocytes and also during the aging of RBCs, The statistical significance is present only for the increase during RBC ageing, although it is borderline also for the maturation of reticulocytes to young RBCs. The "shorter" and "longer" SM subspecies were found to be asymmetrically distributed between the two leaflets. About 70% of those in the inner leaflet, which contains ≈20% of total SM, are of the "shorter" type, while ≈70% of the "longer" SM subclasses are located in the outer leaflet, which contains ≈80% of all SM[14]. According to the cited article, the asymmetric distribution involves only SM subclasses with acyl chains of different length, independently from the degree of unsaturation[14]. Interestingly our results show that during maturation of reticulocytes and especially during ageing of RBCs also the SM subclasses that differ for degree of unsaturation undergo changes, with a slight increase of the more saturated species (**right**). EY, EM, EO, n=4; RM, n=3; RY, n=2. For RY, where only two samples were available, Student's t test could not be performed.

### FIGURE S11

| PS species (mol% of all PS) | RY | | RM | | EY | | EM | | EO | |
|---|---|---|---|---|---|---|---|---|---|---|
| | Mean | * (n=2) | Mean | ± SD (n=3) | Mean | ± SD (n=4) | Mean | ± SD (n=4) | Mean | ± SD (n=4) |
| **18:0-20:4 = 38:4** | 70.6 | ± 8.6 | 74.0 | ± 11.0 | 82.4 | ± 9.4 | 82.7 | ± 7.7 | 84.3 | ± 5.7 |
| **18:0-22:6 = 40:6** | 29.4 | ± 8.6 | 26.0 | ± 11.0 | 17.6 | ± 9.4 | 17.3 | ± 7.7 | 15.7 | ± 5.7 |

*) difference between the mean and each of the two values.

| paired t test (All samples n=4 except RM: n=3 and RY: n=2) | | 38:4 | 40:6 |
|---|---|---|---|
| RM vs. EY | p= | 0.0468 | 0.0468 |
| RM vs. EM | p= | 0.0573 | 0.0573 |
| RM vs. EO | p= | 0.1165 | 0.1165 |
| EY vs. EM | p= | 0.7609 | 0.7609 |
| EY vs. EO | p= | 0.3877 | 0.3877 |
| EM vs. EO | p= | 0.2488 | 0.2488 |

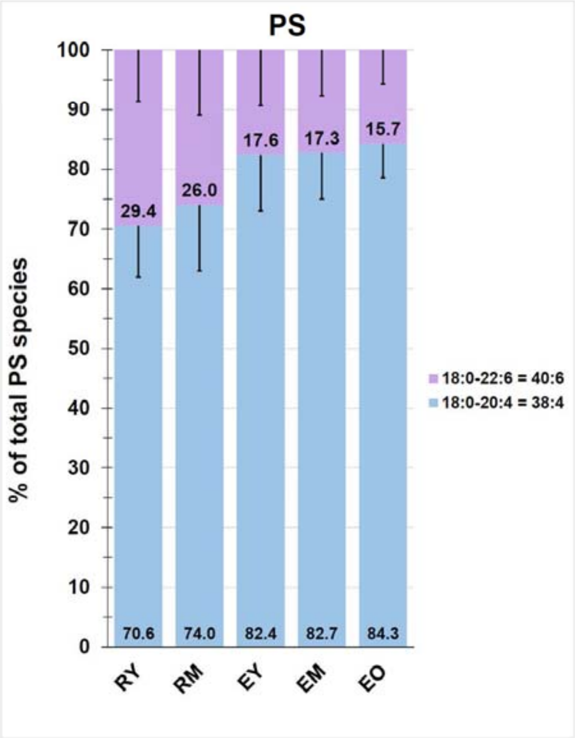

FIGURE S12

| | | paired t test (All samples n=4 except RM: n=3 and RY: n=2) | | | | | | | | | | | |
|---|---|---|---|---|---|---|---|---|---|---|---|---|---|
| | | 12:0 | 13:0 | 14:1 | 16:0 | 17:0 | 18:0 | 18:1 | 18:2 | 20:3 | 20:4 | 21:0 | 22:6 |
| RM *vs.* EY | p= | 0.1690 | 0.2943 | 0.3585 | 0.6148 | 0.8853 | 0.0719 | 0.5037 | 0.2182 | 0.4377 | 0.6516 | 0.2550 | 0.5591 |
| Rm *vs.* EM | p= | 0.3197 | 0.4497 | 0.2780 | 0.0410 | 0.1482 | 0.5146 | 0.0698 | 0.0057 | 0.1380 | 0.6702 | 0.3396 | 0.2645 |
| RM *vs.* EO | p= | 0.0587 | 0.0388 | 0.0505 | 0.0554 | 0.1970 | 0.8484 | 0.1888 | 0.0507 | 0.6625 | 0.0305 | 0.0565 | 0.1902 |
| EY *vs.* EM | p= | 0.5691 | 0.6188 | 0.7431 | 0.2199 | 0.7926 | 0.0411 | 0.2807 | 0.9913 | 0.2583 | 0.8094 | 0.6944 | 0.8160 |
| EY *vs.* EO | p= | 0.8939 | 0.3371 | 0.3804 | 0.2756 | 0.4142 | 0.3408 | 0.5661 | 0.8003 | 0.2794 | 0.1940 | 0.3925 | 0.4691 |
| EM vs. EO | p= | 0.1990 | 0.0407 | 0.0074 | 0.3413 | 0.0848 | 0.3145 | 0.9805 | 0.5324 | 0.3519 | 0.2016 | 0.0736 | 0.4890 |

| paired t test<br>(LPC 12:0 to LPC 18:1) *vs.* (LPC 18:2 to LPC 22:6) | | |
|---|---|---|
| RM *vs.* EY | p= | 0.0258 |
| Rm *vs.* EM | p= | 0.0712 |
| RM *vs.* EO | p= | 0.0193 |
| EY *vs.* EM | p= | 0.8811 |
| EY *vs.* EO | p= | 0.8912 |
| EM vs. EO | p= | 0.8332 |

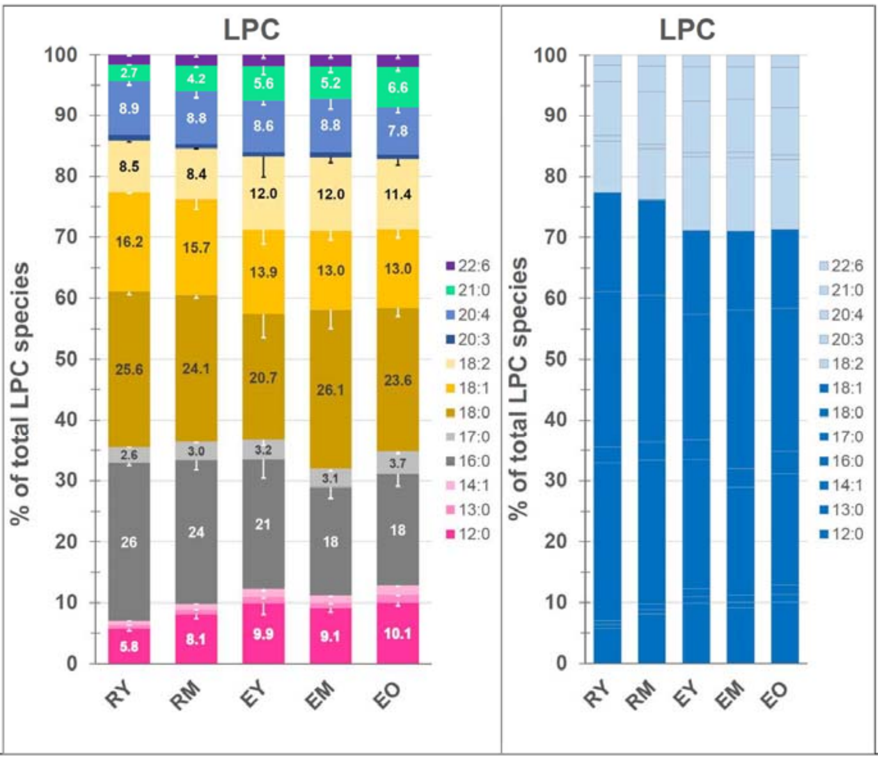

FIGURE S13

| PI species (mol% of all PI) | RY | RM | EY | EM | EO |
|---|---|---|---|---|---|
| | Mean * (n=2) | Mean ± SD (n=3) | Mean ± SD (n=4) | Mean ± SD (n=4) | Mean ± SD (n=4) |
| **16:0-20:4 = 36:4** | 2.3 ± 0.0 | 3.2 ± 0.3 | 7.4 ± 2.1 | 12.6 ± 2.2 | 13.0 ± 3.0 |
| **38:3** | 6.3 ± 2.0 | 6.8 ± 1.9 | 6.3 ± 1.1 | 5.4 ± 0.9 | 4.9 ± 1.3 |
| **18:0-20:4 = 38:4** | 91.4 ± 2.0 | 90.0 ± 2.0 | 86.3 ± 3.0 | 82.0 ± 2.7 | 82.0 ± 4.3 |

*) difference between the mean and each of the two values.

| paired t test (All samples n=4 except RM: n=3 and RY: n=2) | | 16:0-20:4 = 36:4 | 38:3 | 18:0-20:4 = 38:4 |
|---|---|---|---|---|
| **RM *vs.* EY** | p= | 0.1020 | 0.7406 | 0.0563 |
| **Rm *vs.* EM** | p= | 0.0132 | 0.0859 | 0.0122 |
| **RM *vs.* EO** | p= | 0.0347 | 0.1331 | 0.0642 |
| **EY *vs.* EM** | p= | 0.0175 | 0.1236 | 0.0139 |
| **EY *vs.* EO** | p= | 0.0146 | 0.0647 | 0.0534 |
| **EM vs. EO** | p= | 0.5246 | 0.3364 | 0.9453 |

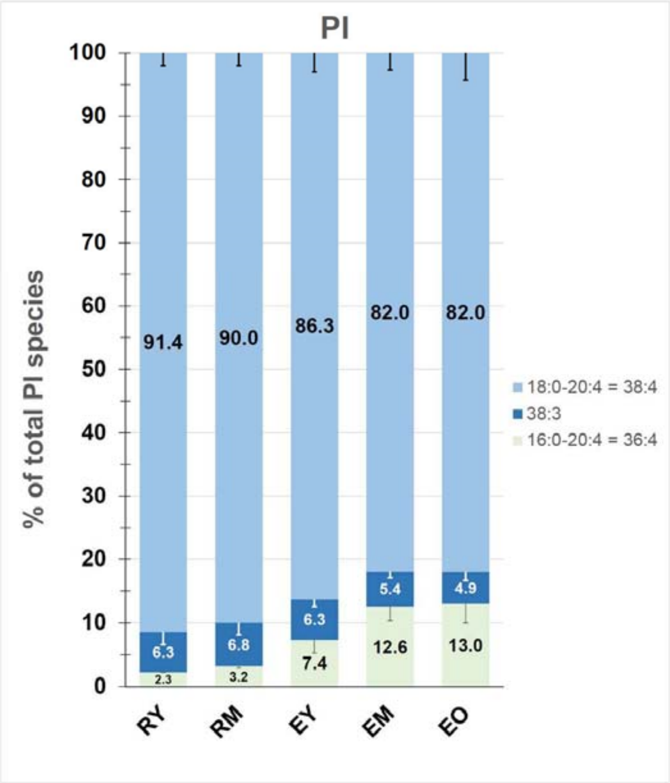

**Table S3. Ratios of palmitoyl-oleoyl to palmitoyl-linoleoyl phosphatidylcholine in the various populations of cells analysed**

| (PC16:0/18:1)/(PC16:0/18:2) | Mean | | SD | C.V.(%) | paired t test (n=4; n=3 for RM | |
|---|---|---|---|---|---|---|
| **RM** | **2.53** | ± | 0.34 | 13.3 | RM *vs.* EY, p= | 0.0176 |
| **EY** | **1.19** | ± | 0.13 | 11.1 | RM *vs.* EM, p= | 0.0120 |
| **EM** | **0.95** | ± | 0.05 | 5.3 | RM *vs.* EO, p= | 0.0115 |
| **EO** | **0.93** | ± | 0.06 | 6.1 | EY *vs.* EM, p= | 0.0206 |
| | | | | | EY *vs.* EO, p= | 0.0210 |
| | | | | | EM *vs.* EO, p= | 0.0898 |

**Table S4. Phospholipid distribution in RBCs from different mammalian species**

| | SM | PC | SM+PC | PE | PS | PE+PS | OTHER |
|---|---|---|---|---|---|---|---|
| RAT | 12.8 | 47.5 | 60.3 | 21.5 | 10.8 | 32.3 | 7.4 |
| DOG | 10.8 | 46.9 | 57.7 | 22.4 | 15.4 | 37.8 | 4.5 |
| HORSE | 13.5 | 42.4 | 55.9 | 24.3 | 18.0 | 42.3 | 1.8 |
| GUINEA PIG | 11.1 | 41.1 | 52.2 | 24.6 | 16.8 | 41.4 | 6.4 |
| RABBIT | 19.0 | 33.9 | 52.9 | 31.9 | 12.2 | 44.1 | 3.0 |
| CAT | 26.1 | 30.5 | 56.6 | 22.2 | 13.2 | 35.4 | 8.0 |
| MAN | 25.8 | 28.3 | 54.1 | 26.7 | 12.7 | 39.4 | 3.9 |
| PIG | 26.5 | 23.3 | 49.8 | 29.7 | 17.8 | 47.5 | 2.7 |
| GOAT | 45.9 | 0.0 | 45.9 | 27.9 | 20.8 | 48.7 | 5.4 |
| COW | 46.2 | 0.0 | 46.2 | 29.1 | 19.3 | 48.4 | 5.4 |
| SHEEP | 51.0 | 0.0 | 51.0 | 26.2 | 14.1 | 40.3 | 8.7 |
| MEAN | 26.2 | 26.7 | 53.0 | 26.0 | 15.6 | 41.6 | 5.2 |
| SD | 15.0 | 18.7 | 4.6 | 3.4 | 3.2 | 5.3 | 2.3 |
| C.V.% | **57.3** | **70.2** | **8.6** | **13.0** | **20.7** | **12.8** | **43.5** |

**Data elaborated from[15].**

FIGURE S14

| p values for the paired t test (all samples n=4 except RM: n=3 and RY: n=2) | | |
|---|---|---|
| | LINOLEIC ACID | ARACHIDONIC ACID |
| RM vs. EY | 0.02477 | 0.21196 |
| RM vs. EM | 0.01064 | 0.01927 |
| RM vs. EO | 0.01055 | 0.01228 |
| EY vs. EM | 0.01253 | 0.06125 |
| EY vs. EO | 0.00947 | 0.00475 |
| EM vs. EO | 0.02435 | 0.03071 |

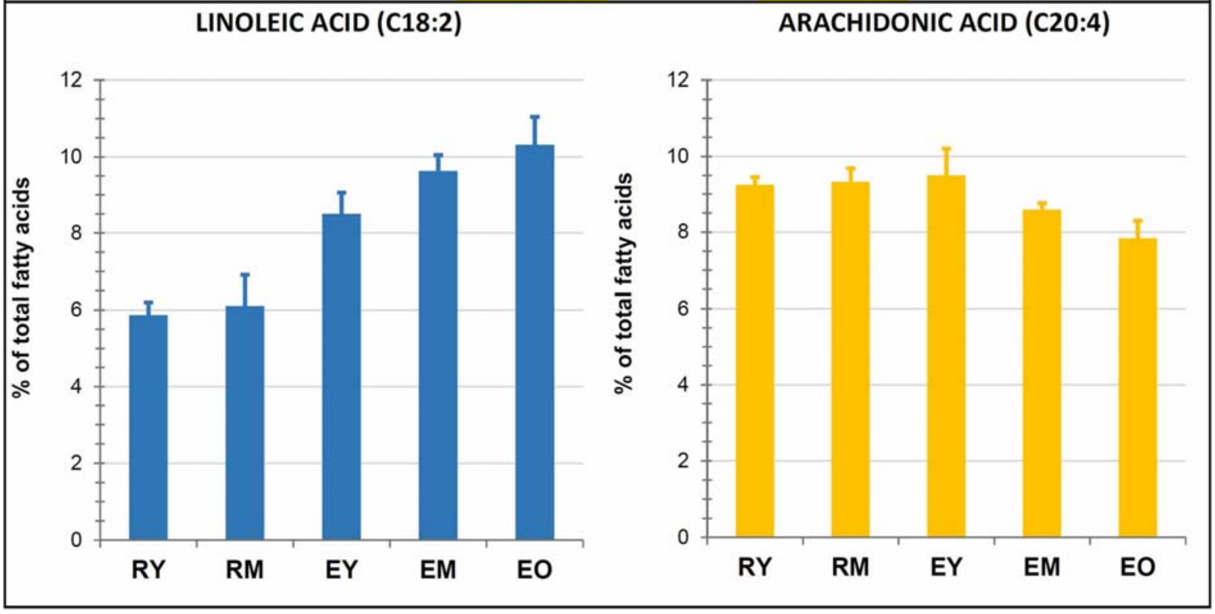


**Figure S14.** The relative amounts of linoleic acid and arachidonic acid with respect to total fatty acids were computed from data on the subclass composition of the individual phospholipids quantified in the study.

#### SUPPLEMENTARY DISCUSSION

Until now, the characterization of terminal erythropoiesis has primarily focused on the progressive simplification of cellular composition, metabolism, and size. Previous studies from transcriptomics and proteomics have indicated a cessation of transcription for most genes encoding RBC proteins early in the process, with the majority of protein species in the membrane and cytosol disappearing as reticulocytes transition into RBCs[13,16-25]. We have uncovered significant compositional changes in the lipid components of the plasma membrane throughout this process. The heterogeneity of young red cells is a key aspect explored in this study.

Unique to our research is the isolation of two distinct subpopulations of circulating reticulocytes: young RY and older RM. Notably, young RY exhibit markedly higher levels of CD71 in the membrane compared to older RM, with a disparity of 20-30 times. Estimating absolute cell age differences based on CD71 content proves challenging due to the unknown kinetics of TfR decay in normal circulating reticulocytes. Moreover, precise quantification of TfR presents complexities and has yielded conflicting findings in previous research[26]. Nonetheless, given that circulating reticulocytes shed all CD71 and RNA within 1-2 days in vivo, it is reasonable to infer that RY represent the first 12-24 hours post-egress from the bone marrow, while RM represent the subsequent 24-48 hours of circulation. Importantly, these reticulocytes were "normal," devoid of contamination by "stress" or "shift" reticulocytes[27], as well as by mature RBCs, distinguishing them from previous studies.

Human RBCs have a potential lifespan of 120 days, equivalent to approximately 17 weeks[28]. Under normal physiological conditions, approximately 0.83% of new reticulocytes enter circulation daily, while an equal number of old RBCs are cleared[29]. The subset of blood cells identified here as EY, is indicative of cells in their first week of life, amounting to around 5% of total blood cells. Conversely, EO cells, representing approximately the 2% densest circulating RBCs, correspond to cells in their 17$^{th}$ week of life. The transition in membrane lipid composition from reticulocytes to EY thus requires at least one week to manifest. Furthermore, EY continue to undergo significant changes, suggesting that it may take several additional days for their membrane to attain the compositional and mechanical stability characteristic of mature RBCs. Consequently, the traditional notion of reticulocyte maturation occurring within 1-2 days in vivo should be re-evaluated; this timeframe primarily reflects the disappearance of conventional reticulocyte markers rather than the production of a mature RBC.

Linked to this issue is the question of whether peripheral reticulocytes can undergo complete maturation into RBCs in vitro. While this concept has garnered some support, it remains contentious. Many studies have utilized "stress" reticulocytes (i.a.[30-32]), and to a lesser extent, peripheral reticulocytes isolated through various methods[33-35]. The outcomes of these studies varied depending on the specific cell parameter used to assess maturation, such as cytoplasmic RNA or membrane TfR levels, which follow distinct pathways with different kinetics. In vivo, the systemic processing of the cell surface removes TfR more rapidly than endogenous nucleases degrade RNA[36], potentially resulting in the coexistence of CD71-negative and RNA-positive reticulocytes. However, in vitro at 37 °C, TfR levels do not decline spontaneously, unlike RNA levels, leading to the erroneous conclusion that peripheral reticulocytes can mature in vitro when only RNA content is considered. Studies adopting TfR decline as a marker for in vitro maturation either did not directly monitor reticulocytes during incubation[33] or utilized stress reticulocytes or those from mice infected with Friend leukemia virus[31,32]. Consequently, it is reasonable to affirm that peripheral circulating reticulocytes cannot be induced to mature into fully formed RBCs in vitro. This conclusion aligns with the understanding that bone marrow reticulocytes and peripheral reticulocytes represent distinct entities, each undergoing maturation through different, sometimes opposing mechanisms. This contrast becomes particularly evident when considering the reduction in membrane area observed in both cell types. Marrow reticulocytes spontaneously release a significant amount of membrane in the form of exosomes, which lack Band 3 and are enriched in sphingolipids and cholesterol[1]. On the other hand, peripheral reticulocytes lose membrane regions that are deficient in sphingolipids and cholesterol but contain Band 3 and spectrin (see Supplementary material, **Figure 1G**). This process necessitates the circulation of the cell within the vascular system.

Even more contentious is the issue as to whether circulating reticulocytes can still synthesize Hb. Experiments suggesting that these cells can produce up to 20% of Hb are likely flawed due to analytical artefacts, potentially stemming from the operation of automated hematology analyzers beyond specified parameters[35]. Moreover, even among the most immature stress reticulocytes observed in experiments involving phenylhydrazine-treated rabbits, only a third of the cells exhibit activity in synthesizing proteins[37]. Hence, a more conservative perspective suggests that no significant alterations in Hb content occur during the maturation of peripheral reticulocytes[38], neither through synthesis nor through loss via vesicles. Conversely, Hb concentration increases due to the decrease in cell volume that accompany reticulocyte maturation[8].

**Size and shape**. While bone marrow reticulocytes shed membrane area through exosome release via a complex membrane trafficking process[39], peripheral reticulocytes lack the molecular machinery to continue reducing surface area through the same mechanism. They may be processed by macrophages with the "pinching" of selected portions of the membrane in the form of vesicles. Volume decrease may be based on the same membrane transport systems in reticulocytes (both marrow and peripheral) and RBCs. In fact, RBCs experience a continual decrease in size and volume until the final day of their circulatory lifespan[40]. Volume regulatory mechanism may involve mechanosensitive channels and loss of potassium through the Gardos channel[41]. In reticulocytes the $K^+/Cl^-$-cotransport, whose activity could still be triggered in young RBCs, to later disappear in mature RBCs[10,42,43], may also be involved. The modulation of these mechanisms during erythroid development[42], including the role of membrane lipids and their remodelling, remains unknown but may be crucial for achieving the necessary cell shrinkage.

It goes perhaps against common belief that the vast majority of circulating reticulocytes appear, as in our hands, as normal biconcave discocytes, with a few misshaped cells of the knizocyte and codocyte (target-cell) type[44], more abundant among the younger reticulocytes (**Figure 2**), which are commonly found among RBCs with excess membrane surface area[45]. Therefore, previous characterizations of reticulocytes as roundish, irregularly shaped, multi-lobular cells, dealt with samples containing significant amounts, perhaps 100%, of marrow reticulocytes[38]. More careful approaches reveal, as in the present study, that circulating reticulocytes are morphologically indistinguishable from mature RBCs[31,46,47].
